# Unraveling patterns of disrupted gene expression across a complex tissue

**DOI:** 10.1101/2021.07.08.451646

**Authors:** Kelsie E. Hunnicutt, Jeffrey M. Good, Erica L. Larson

## Abstract

Whole tissue RNASeq is the standard approach for studying gene expression divergence in evolutionary biology and provides a snapshot of the comprehensive transcriptome for a given tissue. However, whole tissues consist of diverse cell types differing in expression profiles, and the cellular composition of these tissues can evolve across species. Here, we investigate the effects of different cellular composition on whole tissue expression profiles. We compared gene expression from whole testes and enriched spermatogenesis populations in two species of house mice, *Mus musculus musculus* and *M. m. domesticus*, and their sterile and fertile F1 hybrids, which differ in both cellular composition and regulatory dynamics. We found that cellular composition differences skewed expression profiles and differential gene expression in whole testes samples. Importantly, both approaches were able to detect large-scale patterns such as disrupted X chromosome expression although whole testes sampling resulted in decreased power to detect differentially expressed genes. We encourage researchers to account for histology in RNASeq and consider methods that reduce sample complexity whenever feasible. Ultimately, we show that differences in cellular composition between tissues can modify expression profiles, potentially altering inferred gene ontological processes, insights into gene network evolution, and processes governing gene expression evolution.

## INTRODUCTION

A single genome acts as the blueprint for all of the diverse cell types that comprise a eukaryotic organism. This diversity of cellular function is achieved through the expression of individual genes orchestrated by large, layered regulatory networks (Davidson and Erwin 2006; Wittkopp 2007). Often it is through gene expression that changes to the genome are connected to higher level organismal phenotypes of primary interest, and the evolution of gene expression itself can profoundly influence a species’ evolutionary trajectory (King and Wilson 1975; Carroll 2008; Stern and Orgogozo 2008). Gene expression is not a static biochemical phenotype – it is an amalgamation of expression profiles of individual cell types as genes are turned on and off across organismal space and developmental time. Bulk RNASeq of whole tissues allows us to investigate these dynamics in non-model systems with minimal genomic resources and is affordable and tractable for field-based studies (Alvarez et al. 2015). However, evolutionarily important phenotypes often manifest in complex heterogenous tissues, such as sterility in reproductive organs (Turner *et al*. 2012; Suzuki and Nachman 2015), behavioral changes in neurological tissue (Sato *et al*. 2020), or color patterning across the body (Manceau *et al*. 2011; Poelstra *et al*. 2014). Standard bulk sequencing approaches necessarily collapse the complexity inherent to gene expression in these tissues and implicitly assume equivalent proportions of cell types across different comparisons. But if the relative abundance of cell types differs between contrasts, then we may be unable to distinguish regulatory divergence from differences in cellular composition (Good et al. 2010; Montgomery and Mank 2016). What are the consequences of using a whole tissue approach on expression profiles and how does this impact inferences on evolutionary divergence?

Testes are emblematic of a complex tissue and are central to reproductive divergence and speciation. Testes genes are among the most rapidly evolving at the level of protein sequence (Torgerson et al. 2002; Good and Nachman 2005; Turner et al. 2008; Larson et al. 2016) and gene expression (Brawand *et al*. 2011). Sperm, which are produced by the testes, are among the most morphologically diverse animal cells (Pitnick et al. 2009) and are critical in both prezygotic (*e.g.,* sperm competition) and postzygotic (*e.g.,* hybrid sterility) reproductive barriers between species. Studies of whole testes expression have yielded great insights into the evolution of male reproductive traits (*e.g.,* Catron and Noor 2008; Davis et al. 2015; Mack et al. 2016; Ma et al. 2018; Rafati et al. 2018), but relatively few studies have accounted for the cellular complexity of testes, a factor which we expect to complicate evolutionary inference from whole tissues (Good et al. 2010). Testes are dominated by various stages of developing sperm, primarily postmeiotic cells (∼ 70% in house mice; Bellvé et al. 1977), but also present are mitotic precursors, endothelial cells, support cells (White-Cooper et al. 2009), and even multiple types of sperm in some organisms (Whittington *et al*. 2019). The relative proportion of testes cell types is evolvable and plastic (Ramm and Schärer 2014; Ramm et al. 2014) and can vary across species (Lara *et al*. 2018), mating strategies (Firman *et al*. 2015), age (Ernst *et al*. 2019; Widmayer *et al*. 2020), and social conditions (Snyder 1967). For all these reasons, we might expect the cellular composition of testes to differ – sometimes dramatically – between different species, populations, or experimental contrasts.

The cellular complexity of tissues is often due to the developmental complexity of the phenotypes those tissues produce. In testes, undifferentiated germ cells (spermatogonia) undergo multiple rounds of mitosis then enter meiosis (spermatocytes) where they undergo two rounds of cell division to produce four haploid cells (round spermatids). These cells then undergo dramatic postmeiotic differentiation to produce mature spermatozoa. Each of these stages has a unique gene expression profile (Shima et al. 2004; Green et al. 2018; Hermann et al. 2018) and is subject to different selective pressures (Larson et al. 2018). Spermatogenesis in many animals has an additional layer of developmental complexity in the form of the intricate regulation of the sex chromosomes. During early meiosis in mice, the X chromosome is completely transcriptionally inactivated (meiotic sex chromosome inactivation or MSCI; Handel 2004) and remains repressed for the remainder of spermatogenesis (postmeiotic sex chromosome repression or PSCR; Namekawa et al. 2006). Bulk whole testes sequencing aggregates these diverse developmental stages, limiting our resolution into how the molecular mechanisms underlying phenotypic change act in a developmental context (Larson et al. 2018).

The combination of the cellular heterogeneity and developmental complexity of testes is particularly relevant in understanding the evolution of hybrid male sterility. We expect sterile hybrids to have disrupted testes expression (Mack and Nachman 2017; Morgan *et al*. 2020), but sterile hybrids are also likely to have different testes cell composition compared to fertile mice. For example, some sterile house mouse hybrids have only a fourth as many postmeiotic cells (Schwahn *et al*. 2018). These differences in cell composition alone might cause what looks like differential gene regulation associated with hybridization. This is especially problematic when differences in cell composition correspond to developmental timepoints where hybrid expression is disrupted, such as with the disruption of X chromosome inactivation at MSCI (Good et al. 2010; Bhattacharyya et al. 2013; Campbell et al. 2013; Larson et al. 2017), and it is not clear how patterns of stage-specific disruption in hybrids appear in whole testes where stages exhibiting normal and disrupted X regulation are combined. Evidence for disrupted X chromosome regulation in sterile hybrids varies across taxa (Davis et al. 2015; Rafati et al. 2018; Bredemeyer et al. 2021), but outside of mice, most studies have been restricted to whole testes RNASeq. Although these potentially confounding factors are often acknowledged in whole tissue studies (Good et al. 2010; Turner et al. 2014; Davis et al. 2015; Mugal et al. 2020), no systematic effort has been made to distinguish how differences in cellular composition can be distinguished from underlying regulatory dynamics in hybrids using whole testes samples.

Here, we use two analogous RNASeq datasets of fertile and sterile F1 hybrids from *Mus musculus musculus* and *M. m. domesticus* house mice (Mack *et al*. 2016; Larson *et al*. 2017) as a model to investigate the effects of bulk whole tissue sequencing on divergent gene expression. These subspecies form a hybrid zone in Europe where they produce subfertile hybrid males (Turner et al. 2012). F1 hybrid males from wild-derived strains differ in severity of sterility dependent on the strains and the direction of the cross (Britton-Davidian et al. 2005; Good et al. 2008; Mukaj et al. 2020), with more sterile crosses having greatly disrupted cellular composition and gene expression (Good et al. 2010; Bhattacharyya et al. 2013; Campbell et al. 2013; Turner and Harr 2014; Larson et al. 2017; Schwahn et al. 2018). We use comparisons of fertile and sterile reciprocal F1 hybrids to disentangle the effects of cellular composition and disrupted regulatory processes on divergent gene expression. We first examine which cell types contribute to whole testes expression profiles then test predictions about the effects of cell type abundance on whole testes comparisons. Finally, we assess whether signatures of disrupted gene regulation during specific stages of spermatogenesis are detectable in a whole tissue approach and the consequences of whole tissue sampling on differential gene expression. Collectively, we show that inferences from comparative bulk RNASeq approaches are sensitive to changes in cellular composition in complex tissues and advocate for an increased awareness of histology and tissue morphology during study design of RNASeq in non-model systems to account for such effects.

## MATERIALS AND METHODS

### Mouse strains and datasets

We used gene expression data from two recently published datasets analyzing disrupted hybrid gene expression in whole testes (SRA PRJNA286765; Mack et al. 2016) and enriched cell populations across four stages of spermatogenesis (SRA PRJNA296926; Larson et al. 2017). Both studies sequenced transcriptomes from the same wild-derived inbred strains of the mouse subspecies *M. m. domesticus* and *M. m. musculus*, and their F1 hybrids. For each subspecies, two strains were crossed to generate intraspecific F1s to serve as parental controls, without the effects of inbreeding depression on fertility (Good *et al*. 2008). The *M. m. domesticus* mice were generated by crossing the strains WSB/EiJ and LEWES/EiJ, with LEWES dams for the whole testes dataset and WSB dams for the enriched cell dataset (hereafter *dom*). *M. m. musculus* mice were generated by crossing the strains PWK/PhJ and CZECHII/EiJ with PWK dams for the whole testes dataset and CZECHII dams for the sorted cell dataset (hereafter *mus*). F1 hybrid mice with differing severity of sterility were generated by reciprocally crossing LEWES and PWK; PWK female × LEWES male hybrids are mostly sterile (hereafter *sterile*), LEWES female × PWK male hybrids are mostly fertile (hereafter *fertile*). Mack et al. (2016) produced RNASeq libraries from whole testes for each of the four crosses ((2 parental crosses + 2 hybrid crosses) x 3 replicates per cross, N = 12). Larson et al. (2017) used Fluorescence-Activated Cell Sorting (FACS) to isolate enriched cell populations from four different stages of spermatogenesis: Mitosis: spermatogonia (SP), Meiosis^Before X-Inact.^: leptotene and zygotene spermatocytes (LZ), Meiosis^After X-Inact.^: diplotene spermatocytes (DIP), and Postmeiosis: round spermatids (RS) ((2 parental crosses + 2 hybrid crosses) x 3 replicates per cross x 4 cell types per replicate, N = 48). In both studies, libraries were sequenced on an Illumina HiSeq 2000 (100 bp, PE).

### Read mapping and count estimation

We processed both datasets in parallel. First, we used Trimmomatic v.0.38 (Bolger *et al*. 2014) to trim low quality bases from the first and last 5 bp of each read and bases averaging a Phred score of less than 15 across a 4 bp sliding window. We retained reads with a minimum length of 36 bp (Table S1). To avoid mapping bias, we aligned trimmed reads to published pseudo-reference genomes for *M. m. musculus* and *M. m. domesticus* (Huang et al. 2007) using TopHat v.2.1.1 (Trapnell *et al*. 2009) and retained up to 250 alignments per read for multi-mapped reads. We used Lapels v.1.1.1 to convert alignments to the reference mouse genome coordinates (GRCm38.p6) and merged alignments with suspenders v.0.2.6 (Holt et al. 2013; Huang et al. 2014). We summarized read counts for annotated genes (Ensembl Release 96) using FeatureCounts v.1.4.4 (Liao *et al*. 2014) for read pairs that aligned to the same chromosome (-B and -C). We analyzed the count data with and without multi-mapped reads (-M) and across all annotated genes or protein-coding genes only. We found consistent results using all approaches and here present results using only uniquely mapped reads for all annotated genes, unless otherwise specified. Whole testes samples averaged ∼24 million mapped read pairs per sample while sorted cell populations averaged ∼8 million read pairs.

### Characterizing expression patterns

To investigate how expression differed between both datasets, we defined expressed genes as those with a minimum of one Fragment Per Kilobase of exon per Million mapped reads (FPKM) in at least 3 samples within each dataset. This restricted our analysis to 16,824 genes (12,587 protein-coding) in the whole testes dataset and 21,762 genes (14,284 protein-coding) in the sorted cell dataset. We used R v.4.0.2 for all analyses. We conducted expression analyses using the Bioconductor v.3.11 package edgeR v.3.30.3 (Robinson *et al*. 2010) and normalized the data using the scaling factor method (Anders and Huber 2010).

### Effects of cellular composition on whole testes expression

To first determine which cell types were present and contributing to the expression profiles of both datasets, we tested all sample types for the expression of marker genes known to be specifically expressed in certain cell types. We selected three marker genes from seven testes cell types: spermatogonia, spermatocytes, round spermatids, elongating spermatids, Sertoli cells, epithelial cells, and Leydig cells (Raymond *et al*. 2000; Nguyen *et al*. 2002; Maekawa *et al*. 2004; Li *et al*. 2007; Green *et al*. 2018) as well as marker genes from Hermann et al. (2018). This approach allowed us to assess the purity of sorted cell populations by looking for the expression of non-target cell types in sorted cell populations. We were also able to identify which cell types contributed to the expression profile of whole testes.

Next, we tested the hypothesis that differential expression of stage-specific genes in whole tissues can be caused by differences in the relative abundance of cell types between comparisons—in this case *sterile* and *fertile* F1 hybrids (Fig 1; Good et al. 2010). We defined sets of stage-specific genes using our sorted cell populations of each subspecies (Figs S1A, B; Supplemental File 1). We considered a gene to be specific to a given cell population if its median expression (normalized FPKM) was greater than two times its median expression across all other sorted cell populations (*i.e.,* an induced gene approach as in Kousathanas et al. 2014). We then compared the expression of these stage-specific genes in whole testes of *sterile* and *fertile* hybrids. We did this separately for autosomal and X-linked genes because we expected the forces driving patterns of expression to differ between the two. For autosomal genes, we expected expression to be driven largely by differences in cell composition (*e.g.,* fewer later-stage cell types in *sterile* hybrids should lead to lower expression of stage-specific genes from later stages in *sterile* compared to *fertile* whole testes). In contrast, X chromosome inactivation is disrupted in *sterile* hybrids, which should lead to higher expression of stage-specific genes from later stages in *sterile* whole testes. For autosomal genes, we used one-sided paired Wilcoxon signed-rank tests to test if expression of stage-specific genes from more abundant cell types (Mitosis and Meiosis^Before X-Inact.^) was greater in *sterile* hybrid whole testes and if expression of stage-specific genes from less abundant cell types (Meiosis^After X-Inact.^ and Postmeiosis) was lower in *sterile* hybrid whole testes. Because we did not know whether the effects of differing cellular compositions or misregulation of the X chromosome would be stronger for driving expression patterns of stage-specific X-linked genes in whole testes, we used two-sided Wilcoxon signed-rank tests for X-linked genes.

**Fig. 1.**
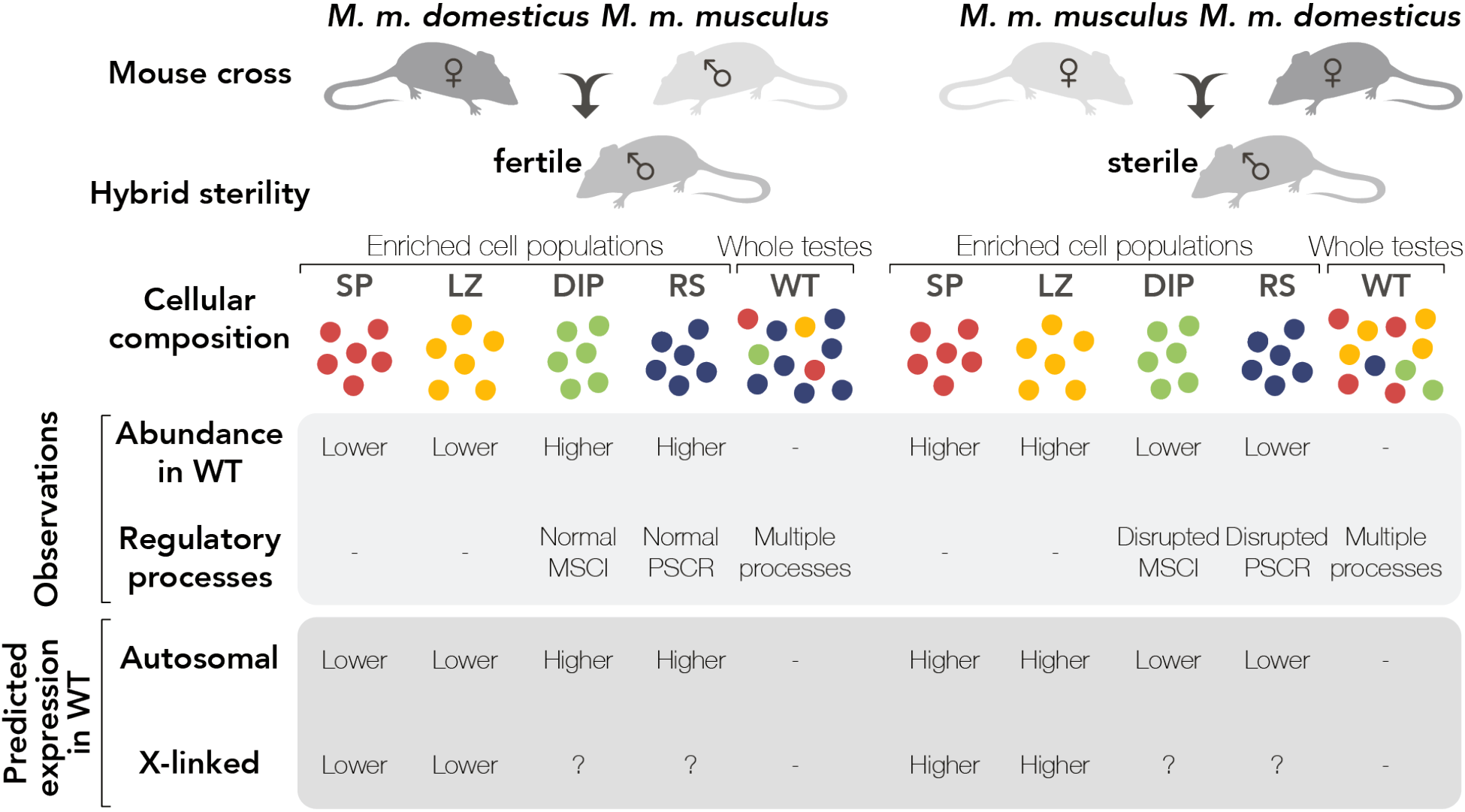
Crossing design, sampling approach, and predicted cell composition and expression differences between reciprocal hybrids. We compared expression patterns using two sampling approaches, enriched cell populations (red = Mitosis (SP), yellow = Meiosis^Before X-Inact.^ (LZ), green = Meiosis^After X-Inact.^ (DIP), blue = Postmeiosis (RS)) and Whole Testes (WT). Whole testes are susceptible to changes in cellular composition between hybrids while enriched cell populations should be buffered from these effects. Relative expression of autosomal stage-specific genes from each enriched cell population in whole testes samples is predicted to track changes in cellular composition between *sterile* and *fertile* mice. Relative expression of X-linked stage-specific genes from enriched cell populations is predicted to be influenced by both changes in cellular composition and expected regulatory processes operating in those cell types.

To look for additional signatures of disrupted X-linked gene expression in both sampling approaches, we first used one-sided Wilcoxon signed-rank tests to compare expression of X-linked genes in *sterile* compared to *fertile* hybrids for each sample type where disrupted X-linked expression was expected (*i.e.,* Meiosis^After X-Inact.^, Postmeiosis, and Whole Testes). Then, we defined sets of “detected” genes for each sample type as those expressed above an FPKM threshold within a replicate (FPKM > 0 – 4) and ran one-way ANOVAs on the number of detected X-linked genes in each cross within a sample type and conducted posthoc Tukey’s tests. Note, because there are only three replicates per sample type, these data inevitability violate distribution assumptions in both parametric and non-parametric tests, and differences among treatments should be considered largely qualitative.

### Differential expression analysis

We conducted differential expression analysis between *sterile* and *fertile* hybrids for all five sample types in edgeR. We fit each dataset (whole testes and sorted cells separately) with negative binomial generalized linear models with Cox-Reid tagwise dispersion estimates (McCarthy *et al*. 2012) and adjusted *P-*values to a false discovery rate (FDR) of 5% (Benjamini and Hochberg 1995). We quantified the biological coefficient of variation (BCV) of parental samples and hybrid samples combined and separately for each dataset. The BCV is the square root of the dispersion parameter from the negative binomial model and represents variation in gene expression among replicates (McCarthy *et al*. 2012). We used two bootstrapping approaches to determine whether BCVs differed across datasets. First, we subsampled raw count files for 10000 genes across 100 replicates and recalculated the BCV for four groups: hybrid whole testes, parent whole testes, hybrid sorted cells, and parent sorted cells. Second, we dropped one individual per group and recalculated the BCV for every set of n-1 individuals. For both approaches, we estimated 99% confidence intervals for the bootstrap BCV estimates from each group and approach (reported as CI_1_ and CI_2_, respectively).

We contrasted expression between *sterile* and *fertile* hybrids so that a positive log fold-change (logFC) indicated over-expression in *sterile* males. For all pairwise comparisons of sample types, we assessed the number of genes overlapping between both sets of differentially expressed (DE) genes and the number of DE genes unique to each sample type in the comparison. We also calculated whether the direction of fold change for a particular DE gene switched between sample types (*e.g.,* an up-regulated DE gene in *sterile* whole testes that was a down-regulated DE gene in any of the *sterile* sorted cell populations). We extended this analysis comparing the direction of DE genes between sample types to parental samples, contrasting expression between *mus* and *dom* parents so that a positive logFC indicated over-expression in *mus* males. We tested for enrichment of specific chromosomes for DE genes between hybrids for each sample type using hypergeometric tests in R (phyper) and adjusted *P-*values to an FDR of 5% (Benjamini and Hochberg 1995). To reduce false positives, we used only the number of autosomal DE genes as the background in the hypergeometric tests because of the known over-expression of the sex chromosomes in *sterile* hybrids (following Larson et al. 2016).

## RESULTS

### Whole testes showed unique expression patterns

Sample type, not cross, was the main driver of differences in expression profiles between samples. All sorted cell populations and whole testes samples grouped into distinct clusters (Fig 2). Within each sample type, parents formed distinct clusters and hybrids had intermediate expression. *Sterile* and *fertile* hybrids each tended to group more closely together within each sorted cell population, but hybrid crosses were intermixed for whole testes and did not form a distinct cluster.

**Figure 2.**
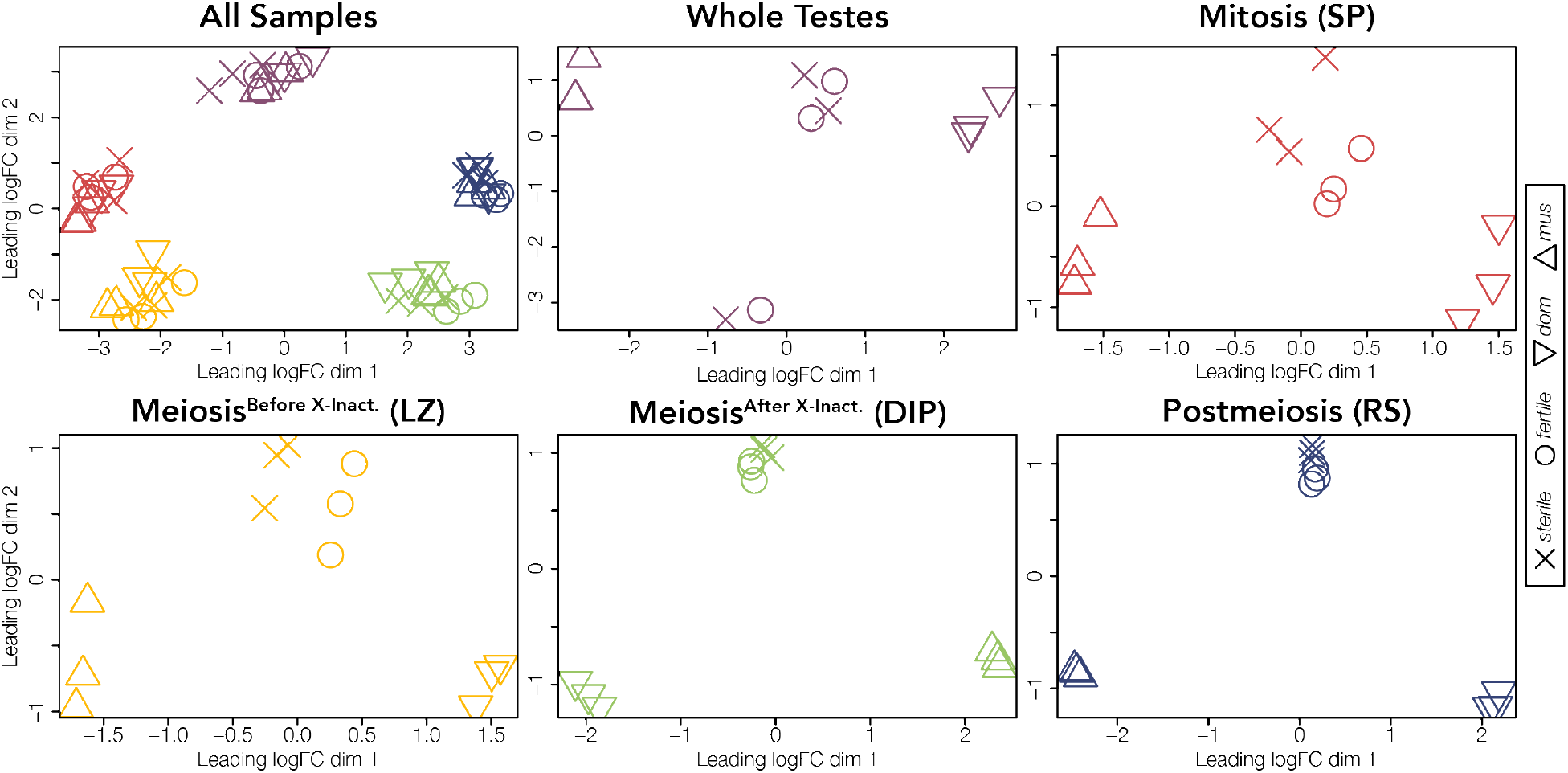
Sample type then cross type drives differences in expression profiles. Multidimensional scaling (MDS) plots of distances among and within sample types for expressed genes across all chromosomes. Distances are calculated as the root-mean-square deviation (Euclidean distance) of log2 fold changes among genes that distinguish each sample. Each cross is indicated by a symbol (*mus* = △, *dom* = ▽, *fertile* = O, and *sterile* = X). Samples are colored by sample type (red = Mitosis, yellow = Meiosis^Before X-Inact.^, green = Meiosis^After X-Inact.^, blue = Postmeiosis, and purple = Whole Testes). The upper left MDS plot includes all sample types and remaining plots show each sample type individually. Each sample type for each cross is represented by three replicates.

Because of the apparent increased variation among whole testes hybrid samples, we next quantified sample variation within both datasets. We measured variation among replicates using the BCV, restricting our analysis to only protein coding genes. Whole testes had greater variation among replicates (BCV = 0.347) compared to sorted cells (BCV = 0.182; Fig S2). Additionally, hybrid whole testes had the greatest variation among replicates (BCV = 0.445; CI_1_ = [0.443,0.445]; CI_2_ = [0.375,0.514]) compared to parent whole testes (BCV = 0.207; CI_1_ = [0.207,0.208]; CI_2_ = [0.161,0.251]), parent sorted cells (BCV = 0.189; CI_1_ = [0.190,0.191]; CI_2_ = [0.185,0.192]), and hybrid sorted cells (BCV = 0.174; CI_1_ = [0.175,0.176]; CI_2_ = [0.171,0.177]; Fig S3-S5). When including all annotated genes in variance calculations, the BCV was still greater in whole testes than in sorted cell populations despite the presence of some lowly expressed and highly variable non-protein coding genes in the sorted cell dataset (Figs S6, S7).

### Whole testes expression patterns are driven by diverse cell composition

We next quantified expression of two panels of marker genes associated with specific testes cell types in fertile reference *mus* and *dom* samples, where gene expression is not expected to be disrupted. This allowed us to assess the purity of sorted cell populations as determined by expression of marker genes from non-target cell types and to ascertain which cell types were contributing to the unique expression patterns observed in whole testes. Our first panel included marker genes associated with spermatagonia (mitosis), spermatocytes (meiosis), round spermatids (postmeiosis), elongating spermatids (postmeiosis), endothelial cells, Sertoli cells (support cells), and Leydig cells (testosterone producing cells), while the second panel included additional cell types (from Hermann et al. 2018; Figs S8, S9). Results from both marker panels were consistent. As expected, sorted cell populations mostly expressed only marker genes characteristic of their target cell type, overall indicating successful FACS enrichment (results for *dom* Figs 3, S8, results for *mus* Figs S9, S10). Mitotic cells showed high expression of spermatogonia markers and limited expression of non-target markers indicating relative cell purity. However, intermediate expression of endothelial and Sertoli markers suggested that the FACS protocol for isolating this cell population may also have captured other somatic cells.

**Figure 3.**
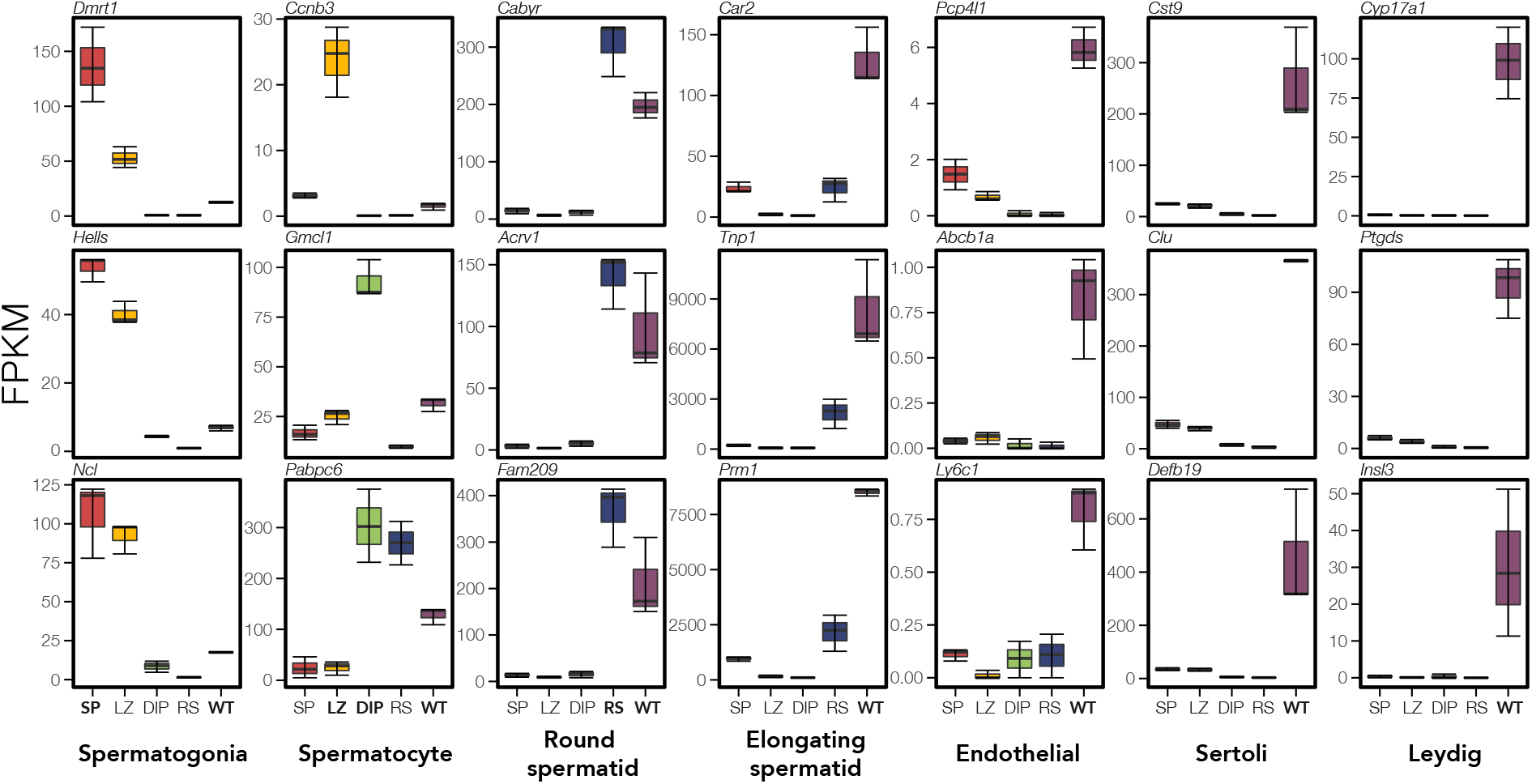
Whole testes show signatures of more diverse cell types than enriched cell populations. Expression of cell-specific marker genes (Green et al. 2018) across each sample type for *dom* reference samples. We quantified expression (FPKM) of three marker genes (rows) associated with testes-specific cell types (columns). Each panel displays marker expression in each sample type (red = Mitosis (SP), yellow = Meiosis^Before X-Inact.^ (LZ), green = Meiosis^After X-Inact.^ (DIP), blue = Postmeiosis (RS), and purple = Whole Testes (WT)). Sample types are bolded in each panel where marker gene expression is expected. Note, *Ccnb3* expression is specific to Meiotic^Before X-Inact.^ cells (Maekawa et al. 2004), and *Gmc1* is specific to Meiotic^After X-Inact.^ cells (Nguyen et al. 2002).

Meiotic^Before X-Inact.^ cells appeared to have some spermatogonia contamination, while Meiotic^After X-Inact.^ cells showed very high purity, expressing only spermatocyte-specific markers. Postmeiotic cells had high expression of round spermatid markers as expected, but also some expression of elongating spermatid markers indicating that FACS may also have captured the developmental transition to these cells.

Whole testes expressed marker genes characteristic of all seven testes cell types, particularly postmeiotic (round and elongating spermatids) and support cell types (endothelial, Sertoli, and Leydig cells) (Fig 3). Additionally, expression patterns on the X chromosome revealed a subset of X-linked genes unique to whole testes samples (Fig 4). These genes were negligibly expressed in each of our sorted cell populations, providing further evidence that additional cell types present in whole testes samples likely contributed to their expression profile. Mitotic (spermatogonia) and meiotic (spermatocyte) markers were also expressed in whole testes but at relatively lower FPKM values, which is consistent with the low relative proportion of these cell types in whole testes (Bellvé *et al*. 1977; Ernst *et al*. 2019). This suggests that early developmental cell types contributed less to whole testes expression profiles, consistent with the hypothesis that the cellular composition of complex tissues can strongly influence relative expression levels (Good et al. 2010).

**Figure 4.**
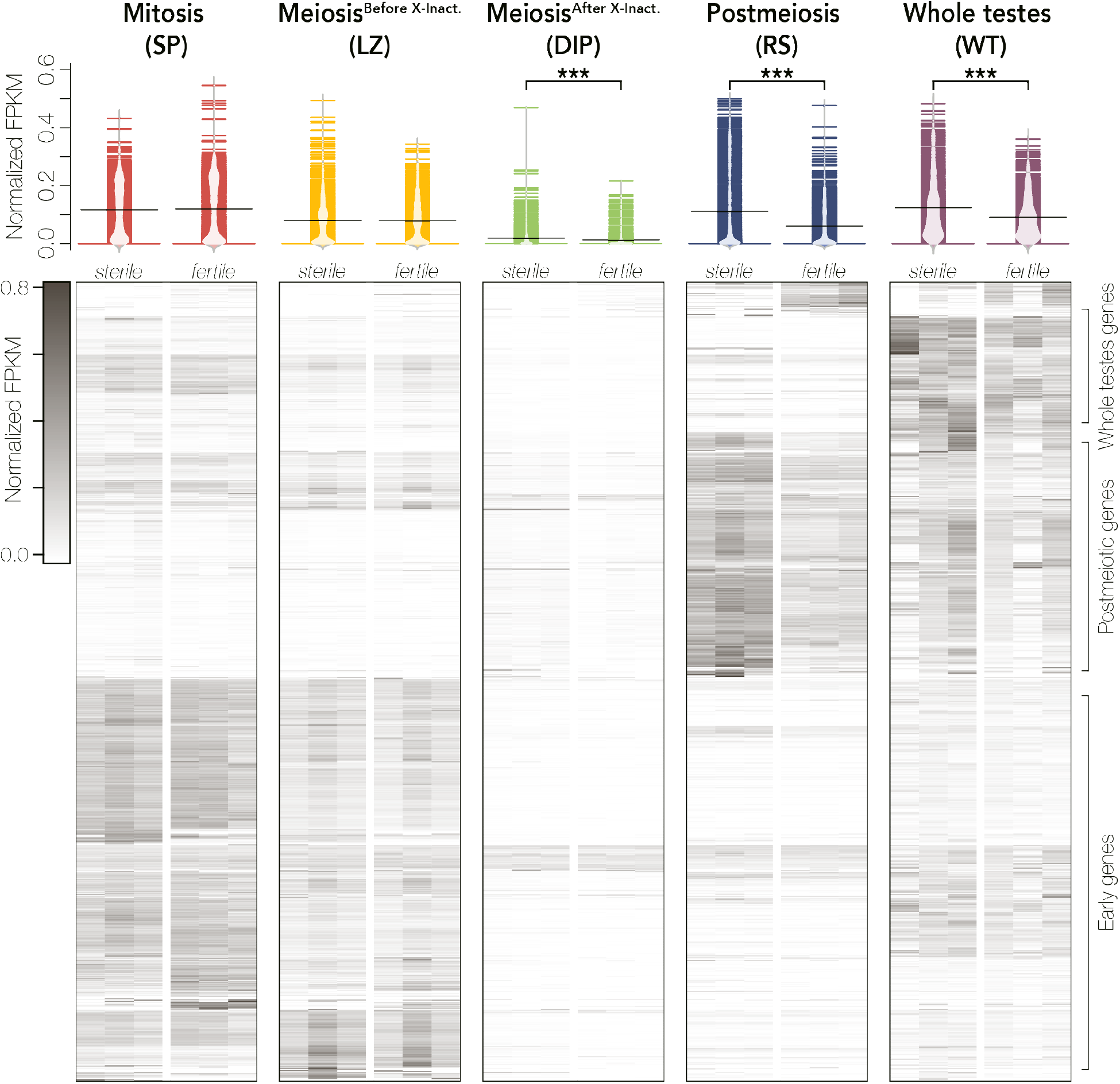
Patterns of X-linked gene expression in *sterile* and *fertile* hybrids differ between sorted cells and whole testes. The upper panel displays expression distributions (as normalized FPKM) across replicates for each sample type across X-linked genes. FPKM values were normalized so that the sum of squares equals one using the R package vegan (Oksanen et al. 2007). Expression distributions are colored by sample type (red = Mitosis, yellow = Meiosis^Before X-Inact.^, green = Meiosis^After X-Inact.^, blue = Postmeiosis, and purple = Whole Testes) and labelled by cross (*sterile* or *fertile* hybrid). These plots were generated with the R package beanplot (Kampstra 2008) and differences in expression were calculated with Wilcoxon signed-rank tests where *** indicates p < 0.001 and * indicates p < 0.05 after FDR correction (Benjamini and Hochberg 1995). The lower panel shows a heatmap of X-linked gene expression plotted as normalized FPKM values that are hierarchically clustered using Euclidean distance. Each row represents a gene with darker colors indicating higher expression. The heatmap was generated with the R package ComplexHeatmap v.2.3.2 (Gu *et al*. 2016).

### Both changes in cellular composition of whole testes and regulatory divergence contribute to expression differences in hybrids

We further tested whether changes in cellular composition of complex tissues influences relative expression levels between contrasts. Indeed, we found that differences in whole testes cell composition between *sterile* and *fertile* hybrids appears to be a large driver of differences in relative expression of stage-specific genes (Fig 5). In *fertile* hybrids, whole testes are largely composed of late spermatogensis cell types. In *sterile* hybrids, there is a disruption in development immediately before normal MSCI, which triggers an apoptotic cascade and decreases downstream meiotic and postmeiotic cell abundance (Schwahn *et al*. 2018). Based on these histological predictions, we expected stage-specific genes from pre-X chromosome inactivation stages (Mitosis and Meiosis^Before X-Inact.^) to appear over-expressed in *sterile* hybrids and stage-specific genes from post-X chromosome inactivation stages (Meiosis^After X-Inact.^ and Postmeiosis) to appear under-expressed in *sterile* hybrids. Consistent with this, in whole testes, autosomal Mitotic- and Meiotic^Before X-Inact.^-specific genes had higher expression in *sterile* hybrids (one-sided Wilcoxon Signed-Rank Test; autosomal Mitotic: n = 5307, V = 11247685, p = 0; autosomal Meiotic^Before X-Inact^: n = 4215, V = 7988923, p = 0), while autosomal Meiotic^After X-Inact.^- and Postmeiotic-specific genes had lower expression (one-sided Wilcoxon Signed-Rank Test; autosomal Meiotic^After X-Inact^: n = 4544, V= 2005025, p = 1.46 x 10^-276^ ; autosomal Postmeiotic: n = 7417, V = 4789686, p = 0; Fig 5).

**Figure 5.**
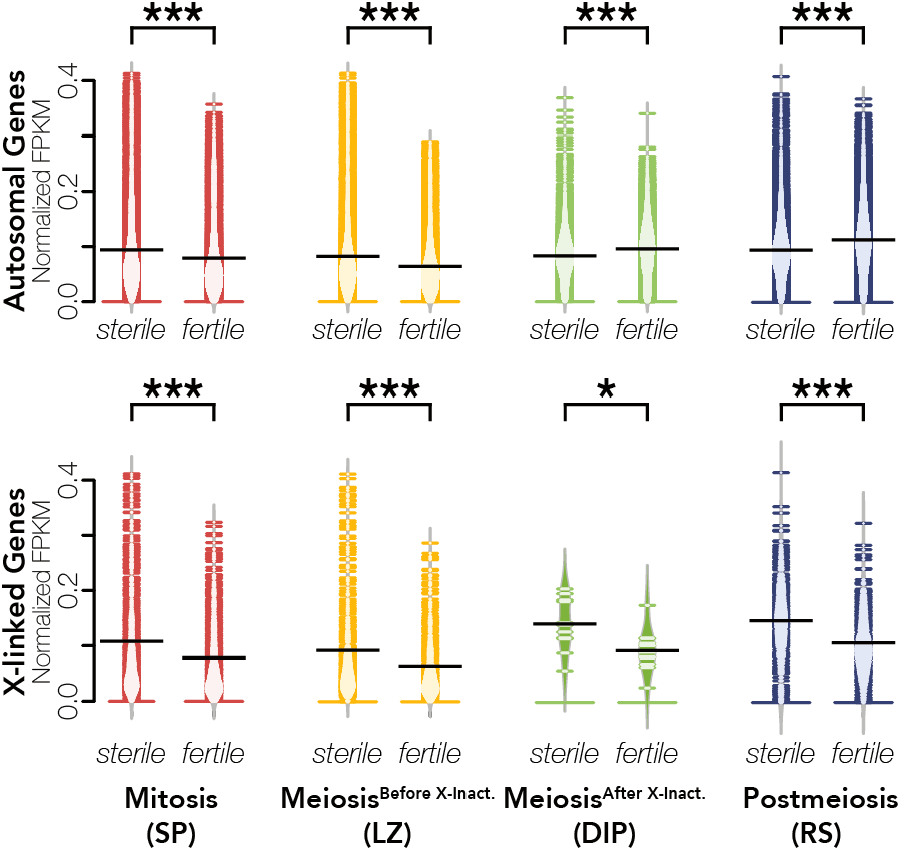
Changes in cellular composition alters expression of stage-specific genes in whole testes samples. For each sorted cell population, we defined a set of stage-specific genes and compared their expression in whole testes of *sterile* and *fertile* hybrids. Mitotic and Meiotic^Before X-Inact.^ cells are present at lower abundances in *sterile* hybrids while Meiotic^After X-Inact.^ and Postmeiotic cells are present at higher abundances (Schwahn et al. 2018). FPKM is normalized so that the sum of squares equals 1 using the R package vegan (Oksanen et al. 2007). Differences in expression were calculated with Wilcoxon signed-rank tests where *** indicates p < 0.001 and * indicates p < 0.05 after FDR correction (Benjamini and Hochberg 1995).

Given the nature of hybrid sterility in house mice (Bhattacharyya et al. 2013), we had different expectations for X-linked genes. The normal regulation of the X chromosome is not disrupted in pre-X inactivation cell types, so differences in cellular composition should drive expression patterns for stage-specific X-linked genes in pre-X inactivation cell types as with autosomal genes. However, the X chromosome is over-expressed in post-X inactivation cell types (Larson et al. 2017), so changes in cellular composition and known regulatory divergence could influence expression patterns of post-X inactivation stage-specific genes in *sterile* whole testes. As we predicted based on cell composition, X-linked Mitotic and Meiotic^Before X-Inact.^ genes still had higher expression in *sterile* hybrids (one-sided Wilcoxon signed-rank test; X-linked Mitotic: n = 465, V = 95946, p = 1.16 x 10^-47^; X-linked Meiotic^Before X-Inact^: n = 361, V = 60492, p = 1.53 x 10^-44^). However, X-linked Meiotic^After X-Inact.^ and Postmeiotic genes also had higher expression (two-sided Wilcoxon Signed-Rank Test; X-linked Meiotic^After X-Inact.^: n = 11, V = 56, p = 0.0420; X-linked Postmeiotic: n = 252, V = 25826, p = 1.59 x 10^-17^), indicating that the disruption of X chromosome inactivation and repression in *sterile* hybrids had a stronger effect on expression patterns than changes in cell composition, despite the lower abundances of these cell types (Schwahn et al. 2018). Together these results indicate that the high proportion of postmeiotic cells in whole testes is a major cause of differences in expression patterns of autosomal and many X-linked genes between *sterile* and *fertile* whole testes samples.

We further investigated the detectability of patterns of disrupted X chromosome regulation in *sterile* hybrids across both sampling approaches and found that whole testes sampling partially masks signatures of X chromosome misexpression. Previous research using sorted cell populations has shown that disruption of MSCI in *sterile* hybrids manifests as over-expression of the X chromosome both in terms of more expressed X-linked genes and higher average X-linked gene expression (Larson et al. 2017). We recovered the expected pattern of higher X-linked gene expression in *sterile* hybrids in both sorted cell populations (one-sided Wilcoxon signed-rank test; X-linked Meiosis^After X-Inact^: n = 896, V = 273584, p = 5.34 x 10^-60^; X-linked Postmeiosis: n = 896, V = 290110, p = 1.18 x 10^-64^; Fig 4) and in whole testes (one-sided Wilcoxon signed-rank test; n = 896, V = 326947, p = 2.40 x 10^-64^; Fig 4). We also found more detected X-linked genes in Meiosis^After X-Inact.^ (F_3,8_ = 13.8, p = 1.58 x 10^-3^; Fig S11A) and Postmeiosis (F_3,8_ = 31.87, p = 8.47 x 10^-05^; Fig S11B), but there was no difference in the number of detected X-linked genes in *sterile* whole testes (F_3,8_ = 0.606, p = 0.629; Fig S11C), regardless of the FPKM threshold used to define detected genes.

### Whole testes sampling reduces power for differential expression inference

The increased variance among replicates and the resulting decreased power in the whole testes dataset also greatly reduced the number of genes considered differentially expressed between *sterile* and *fertile* hybrids in whole testes compared with sorted cell populations (Fig 6; Table S2; Supplemental File 2). Fewer DE genes were detected between hybrids for whole testes samples (DE genes = 83; Table S2) compared to sorted cell populations (Mitotic DE genes = 231, Meiotic^Before X-Inact.^ DE genes = 178, Meiotic^After X-Inact.^ DE genes = 343, and Postmeiotic DE genes = 606). However, both whole testes and sorted cell populations exhibited similar broad patterns of differential expression. In both datasets, more DE genes were upregulated in *sterile* hybrids than were downregulated (Table S2), and we were able to detect enrichment of the X and Y chromosomes for DE genes as previously reported (Larson et al. 2017; Fig S12; Tables S3, S4). In addition, no DE genes between *sterile* and *fertile* hybrids were differentially up- or down- regulated in whole testes samples compared to sorted cell populations (Table S5; Fig S13), although we did find this pattern when comparing DE genes between *mus* and *dom* mice— a small proportion of genes were differentially regulated in whole testes samples compared to sorted cell populations (0.43% - 3.16%; Table S6; Supplemental Files S3, S4).

**Figure 6.**
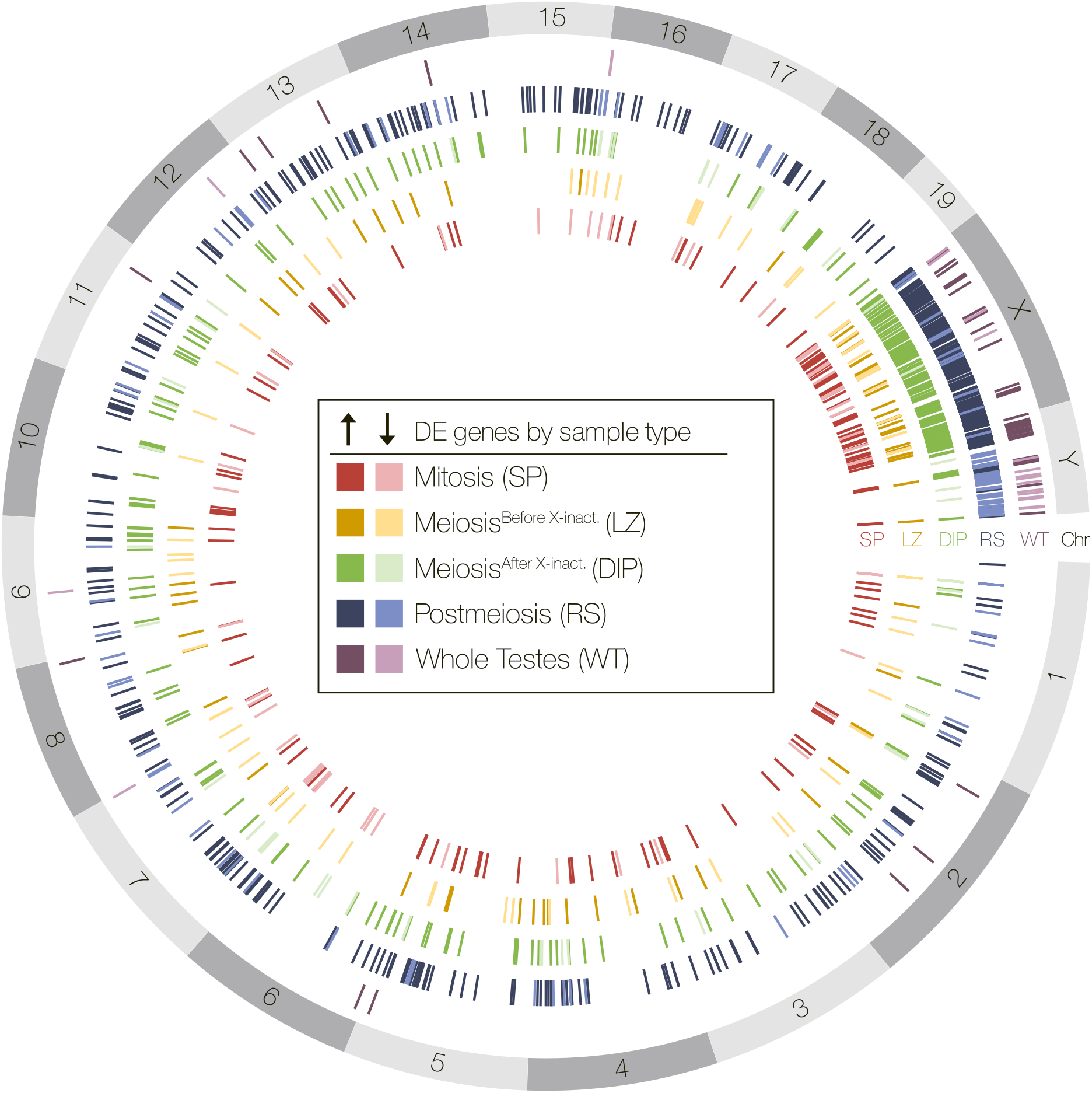
Whole testes and enriched cell populations differed in the number and identity of dfferentially expressed genes. Spatial distribution of differentially expressed (DE) genes across reference mouse genome chromosomes (build GRCm38.p6) between *sterile* and *fertile* hybrids for all five sample types. Darker colors indicate genes up-regulated in *sterile* hybrids and lighter colors indicated genes down-regulated in *sterile* hybrids.

Despite consistent patterns of enrichment of DE genes on the sex chromosomes and direction of expression of DE genes between hybrids, there was very little overlap in DE genes between each sample type. Whole testes samples shared very few genes in common with any of the sorted cell populations (Fig 7). Additionally, there were very few DE genes shared across the different stages of spermatogenesis, although the proportion of DE genes shared between sample types generally increased with stricter fold change cutoffs (Table S7). Sorted cell samples often have large repertoires of genes that were only differentially expressed within one cell type (Fig 7) though there was greater overlap of DE genes between post-X inactivation cell types (Meiosis^After X-Inact.^ and Postmeiosis). In sum, different sampling methodology clearly altered the overall and gene-specific resolution of the regulatory underpinnings of hybrid male sterility.

**Figure 7.**
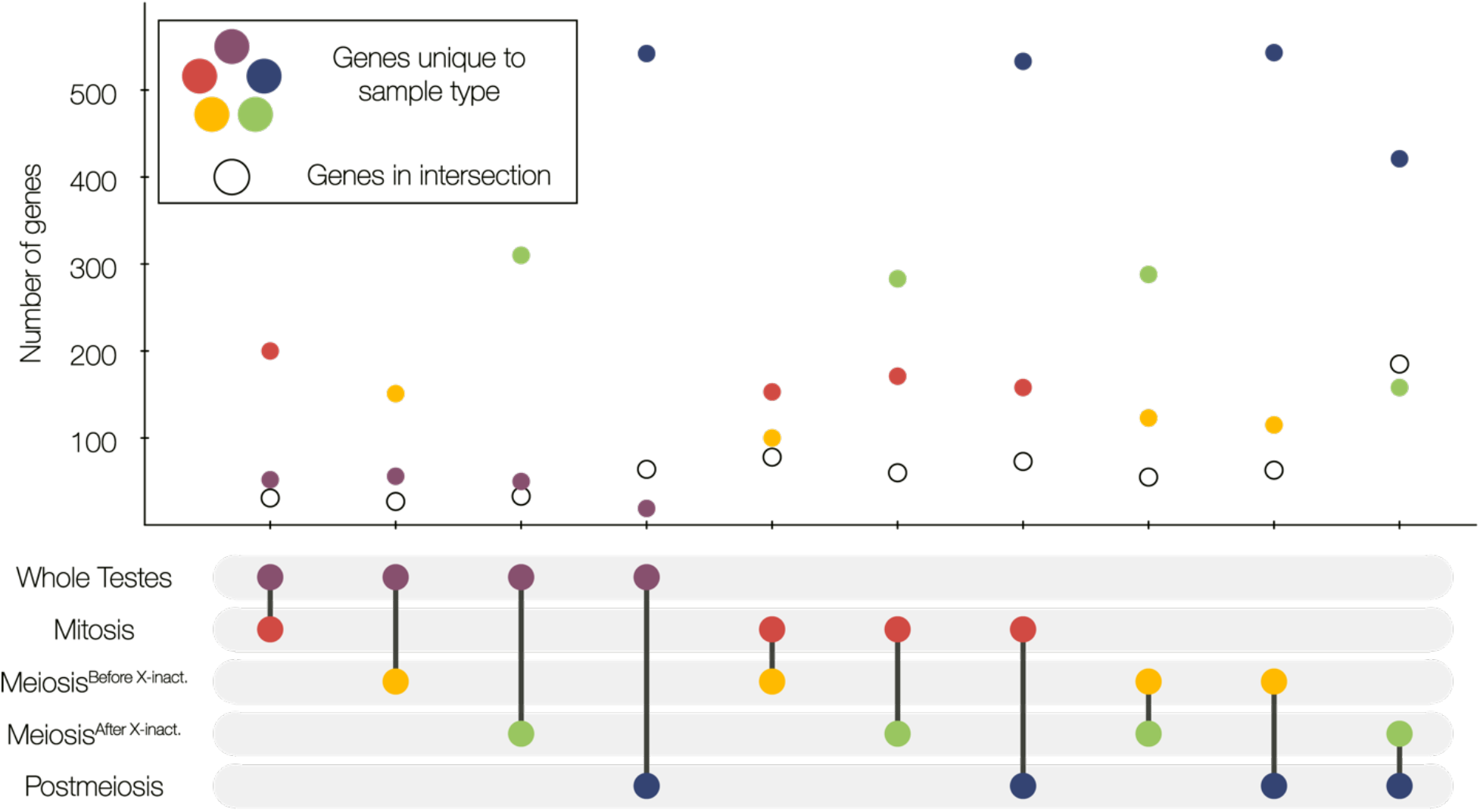
Whole testes and enriched cell populations differed in pairwise comparisons of DE genes between *sterile* and *fertile* hybrids. The sample types in each comparison are indicated by the pair of connected dots in the bottom panel. For each comparison, DE genes common between the two sample types are indicated with a hollow circle and DE genes unique to each sample type in that comparison are colored by sample type (red = Mitosis, yellow = Meiosis^Before X-Inact.^, green = Meiosis^After X-Inact.^, blue = Postmeiosis, and purple = Whole Testes).

## DISCUSSION

### Transcriptomic biases of complex tissues in evolutionary biology

Bulk RNASeq of whole tissues has been the canonical method for characterizing divergent expression in evolutionary biology as it is both cost-effective and tractable for wild populations (Wang et al. 2009; Alvarez et al. 2015; Todd et al. 2016). Here we characterized patterns of expression divergence in *sterile* and *fertile* F1 hybrid house mice that differ in cellular composition using two approaches, whole testes sequencing and isolation of enriched cell populations across different stages of spermatogenesis. We demonstrated that bulk RNASeq of this complex tissue strongly reflected the cumulative contributions of diverse cell types and that the relative cell type proportions in *sterile* and *fertile* hybrids influenced the expression of stage-specific genes. This suggests that differential expression in whole tissues can be due to either cell composition or regulatory divergence, and while these reflect fundamentally different mechanisms, they may be confounded in comparisons between species. This is a critical distinction given that researchers often interpret patterns of gene expression as reflecting per cell changes in transcript levels. This biological interpretation is implicit in models of expression evolution (Rohlfs and Nielsen 2015), which typically assume that cellular composition is stable across species of interest. We must consider the cellular context of divergent gene expression patterns (Montgomery and Mank 2016; Breschi et al. 2017; Buchberger et al. 2019), as the tissues in which these phenotypes occur, such as reproductive organs (Ramm and Schärer 2014), nervous tissues (Carlson et al. 2011; Davidson and Balakrishnan 2016), and plumage (Abolins-Abols et al. 2018; Price-Waldman et al. 2020), may be prone to structural evolution, making them extremely susceptible to confounded mechanisms inherent to whole tissue sampling.

Reproductive tissues are likely to be particularly prone to structural divergence, as cellular composition is expected to evolve in response to selection for increased reproductive success. For example, sperm competition leads to selection for males to increase sperm numbers (Firman et al. 2013, 2018) or the proportion of sperm-producing tissue within the testes (Lüpold et al. 2009). Sperm production can be increased in multiple ways, each of which has different consequences for the cellular architecture of the testis (Schärer *et al*. 2011; Ramm and Schärer 2014). The non-sperm-producing tissue within the testes can also evolve in response to sexual selection. An extreme example are Capybara, which devote ∼30% of their testes to the testosterone-producing Leydig cells (in sharp contrast to other rodents, where Leydig cells comprise only 2.7-5.3% of testes; Costa et al. 2006; Lara et al. 2018). Differences in reproductive investment can also drive apparent expression differences between species. Gene expression divergence between humans and chimpanzees is elevated in testes relative to other tissues, a pattern proposed to reflect positive selection on gene expression levels (Khaitovich et al. 2005, 2006). However, whole testis transcriptomes tend to be more similar between species with similar mating systems and cellular architectures (Brawand et al. 2011; Yapar et al. 2021), which have presumably evolved convergently in response to investment in sperm production. Our results show that in bulk tissues even minor testis cell types (such as Leydig cells and Sertoli cells) contribute to overall expression profiles and suggest that differences in the proportion of any cell type have the potential to strongly modify expression profiles of whole tissues.

### Reducing sample complexity in evolutionary studies of expression divergence

Here we confirm that FACS is an effective way of isolating relatively pure cell types and removing the effects of divergent cellular composition from experimental contrasts (Getun et al. 2011; da Cruz et al. 2016; Larson et al. 2016, 2017; Geisinger et al. 2021; Kopania et al. 2021). There are of course, alternative methods available for bulk cell enrichment, such as gradient centrifugation to separate testes cell types (Shima et al. 2004; Chalmel et al. 2007; Rolland et al. 2009). These approaches are well suited for testes, given the dramatic changes in cell size, DNA content, and chromatin condensation during spermatogenesis (Bellvé 1993; Getun et al. 2011), but mechanical or flow cytometry-based enrichment has also been developed for other complex heterogeneous tissues (e.g., late term placenta; Li et al. 2020). There is the potential for FACS to bias gene expression in enriched cell populations (*i.e.,* by triggering a stress response from nozzle pressure or UV exposure; Box et al. 2020). However, cell sorting procedures appear to have a minimal effect on overall expression profiles, and altered expression is likely consistent across treatments within an experiment and can be mitigated by minimizing the time between cell sorting, RNA extraction, and storage (Box et al. 2020). Beyond the limited and potentially tissue-specific methods of bulk cell enrichment, recent advances in single cell sequencing technology (scRNA-Seq) and spatial transcriptomic methods (*e.g.,* sci-SPACE, Srivatsan et al. 2021) can allow researchers to assay a greater number of cell types across many tissue types without *a priori* identification or labelling (Kiselev et al. 2019). Although both FACS and scRNA-Seq are both powerful approaches for studying gene expression evolution in tissues with cellular composition differences (Kopania et al. 2021; Murat et al. 2021), they are both currently difficult to apply in non-model systems, especially for field-based studies, as they typically require access to flow cytometers and a short timeline for tissue biopsy, cell sorting, and RNA extraction (Getun et al. 2011; Bageritz and Raddi 2019, but see also Wohnhaas et al. 2019; Denisenko et al. 2020). Additionally, scRNA-Seq protocols likely have some of the same sources of biased gene expression as FACS, and further research should be done to determine how different enrichment protocols alter expression inferences.

When cell enrichment protocols are not feasible, alternative methods are available for minimizing developmental or cellular complexity differences between species or experimental contrasts. For example, different stages of sperm development can be isolated by sampling whole testes at different points in early sexual development (Schultz et al. 2003; Shima et al. 2004; Laiho et al. 2013), across annual reproductive cycles (Rolland et al. 2009), or spatially, as in *Drosophila*, where sperm develop in tubular testes, allowing dissection of distinct regions that are enriched for particular cell types (Meiklejohn et al. 2011; Landeen et al. 2016). Furthermore, some developmentally heterogenous samples can be artificially synchronized, for example by shaving hair or plucking feathers and sampling across regrowth timelines (Poelstra et al. 2014, 2015; Ferreira et al. 2017). Microdissection of complex tissues is also a feasible way to minimize the effects of cellular composition on transcriptomic profiles. For example, laser capture microdissection provides a means to rapidly and precisely isolate cellular populations from complex tissues (Emmert-Buck et al. 1996), albeit with the added requirement of highly specialized instrumentation. It is common in behavioral research to dissect out major regions of the brain rather than sampling the whole brain (Khrameeva et al. 2020; Sato et al. 2020). Thus, a chemical or mechanical approach to partitioning complex tissues can provide researchers with a way of minimizing the negative effects associated with bulk RNASeq in their own studies.

Despite the potentially confounding effects of cellular composition and regulatory divergence in whole tissue sampling, a bulk RNASeq approach is appropriate in cases where a cell type of interest is not easily isolated or when researchers wish to capture all developmental stages. For example, Larson et al. (2017) used FACS to isolate only four stages of spermatogenesis, but postzygotic isolation barriers can operate at many different stages of spermatogenesis (Oka et al. 2010; Ishishita et al. 2015; Torgasheva and Borodin 2016; Schwahn et al. 2018; Yoshikawa et al. 2018; Liang and Sharakhov 2019). In these situations, bulk RNASeq can allow researchers to investigate expression differences in hard to obtain cell types. Indeed, some evolutionary inferences may be robust to sampling strategy. The misexpressed genes in hybrids identified by Mack et al. (2016) overlapped substantially with sterility eQTLs identified in wild hybrids from natural hybrid zones of *M. m. musculus* and *M. m. domesticus* populations (Turner and Harr 2014), suggesting that despite the decreased power and susceptibility to artifacts introduced by differences in cellular composition associated with bulk tissue sampling, the genes that are identified are likely genes of large effect and have a high likelihood of being biologically meaningful. For all these reasons, bulk tissue sampling may be an appropriate first step depending on the system and questions being addressed.

It is also possible to use computational approaches, such as *in silico* deconvolution methods to estimate changes in cell type proportions across samples or quantify cell type-specific expression profiles (Shen-Orr and Gaujoux 2013; Avila Cobos et al. 2018; Newman et al. 2019). These methods rely on expression profiles from single-cell data and accurate estimates of cellular proportions (Shen-Orr and Gaujoux 2013; Avila Cobos et al. 2018), which can be challenging to obtain in non-model systems but are likely to become increasingly more accessible as technologies advance. Deconvolution may also be less accurate when the expression of specific genes varies across stages because the net expression of a gene in a whole tissue may differ from its stage-specific expression. While we found that DE genes between *sterile* and *fertile* hybrids had consistent direction of differential expression between our whole testes samples and sorted cell populations, in our comparisons of DE genes between *mus* and *dom* mice, we found DE genes that had the opposite regulation patterns between sample types. Deconvolution methods in studies of hybrid misexpression may also be inherently flawed given that there is often no single “sterile” phenotype (Good et al. 2008; Turner et al. 2012; Larson et al. 2017; Bikchurina et al. 2018) and that the reference expression profiles used for deconvolution may be disrupted in hybrids (Landeen et al. 2016; Morgan et al. 2020; Mugal et al. 2020; Brekke et al. 2021). Given these drawbacks, we advocate that detailed histological analysis of how the phenotype of interest manifests in complex, heterogenous tissues (Oka et al. 2010; Schwahn et al. 2018) should accompany any evolutionary study based on comparative transcriptomic data, so that researchers can mediate biases associated with sampling methodology when designing future studies.

### Power to detect differential expression using bulk RNASeq

The primary analytical goal of most RNASeq studies is to identify DE genes. It is vital that we can accurately determine which genes are differentially expressed because we use these patterns for a myriad of downstream analyses. Accurate assessment should also increase resolution into the genomic basis of phenotypes of interest. We found that bulk RNASeq can hinder differential expression analyses through an increase in replicate variability, potentially masking biologically meaningful changes in gene expression. RNASeq analyses are sensitive to both technical and biological variation (Todd et al. 2016), and studies of outbred wild populations are inherently disadvantaged because of the power lost from increased biological variation (Liu et al. 2014; Todd et al. 2016). The BCV is an estimate of the variation among biological replicates and is correlated with power to detect DE genes. We found that in inbred strains of house mice, whole testes had higher inter-replicate variability in expression than sorted cell populations and levels of variation closer to what would be expected for an outbred wild population (BCVs greater then 0.3; McCarthy et al. 2012; Todd et al. 2016) than for genetically identical model organisms (BCV less than 0.2). We suggest that reporting BCV should become a best-practices standard for all RNASeq studies so that researchers may better understand the nature of biological variation in gene expression across a variety of evolutionary contrasts.

Consistent with the increased BCV in whole testes samples, we found that fewer genes were differentially expressed in whole testes samples than in sorted cell populations and that DE genes in whole testes had little overlap with DE genes in sorted cell populations. However, this overlap proportionally increased with stricter fold change cutoffs, which strongly supports using these cutoffs to decrease the chance of detecting false positive DE genes (as proposed by Montgomery and Mank 2016). The downside to this more conservative approach was that the higher fold change cutoffs likely led to the exclusion of some genes with biologically relevant expression differences.

Ultimately, both whole tissue and cell enrichment-based approaches were able to detect broad-scale patterns of disrupted sex chromosome expression in *sterile* hybrids. In house mice, MSCI is disrupted in *sterile* hybrids (Bhattacharyya et al. 2013; Davies et al. 2016; Gregorova et al. 2018), leading to an over-expression of X-linked genes (Good et al. 2010; Campbell et al. 2013; Turner et al. 2014). Both Mack *et al*. (2016) and Larson *et al*. (2017) found higher expression of genes across the X chromosome in *sterile* hybrids, but our results show that it is more difficult to detect an increased number of expressed X-linked DE genes between *sterile* hybrids and their parents using whole testes sampling. Patterns of X-linked over-expression can also be recovered in whole testes given *a priori* knowledge of stage-specific genes for cell types where the X chromosome should be inactivated or repressed. Of course, approaches relying on orthologous sets of stage-specific genes from other species will be limited to species with close evolutionary relationships to model organisms. A sensitivity of the regulatory mechanisms controlling sex chromosome expression during male meiosis has been proposed to be a major mechanism underlying hybrid sterility (Lifschytz and Lindsley 1972), but so far, genomic evidence for disrupted MSCI and downstream postmeiotic repression in other mammalian taxa is conflicting. In sterile hybrid cats, the X chromosome is misexpressed and MSCI is disrupted (Davis et al. 2015; Bredemeyer et al. 2021), while sterile rabbit hybrids do not support a role of X chromosome misexpression in speciation (Rafati *et al*. 2018). Studies outside of house mice have largely relied on bulk whole testes sequencing (but see Bredemeyer et al. 2021) and understanding if the detected or undetected misexpression of the X is biologically accurate is important for determining the role of disrupted sex chromosome regulation in postzygotic isolation and speciation. Using targeted approaches can give us the developmental perspective needed for contextualizing the origins of reproductive barriers (Cutter and Bundus 2020).

### Conclusions

Here, we demonstrate important consequences of differing cell composition in identifying DE genes in the context of hybrid sterility. We advocate for sampling approaches which allow for developmental perspectives in RNASeq studies, so that we can accurately probe species barriers. These same issues are important for other evolutionary contrasts in complex tissues, and we underscore the importance of considering the cellular and developmental context of complex expression in evolutionary studies. Our results suggest that sampling methodology could influence the biological implications of not only hybrid misexpression in speciation, but also across studies of divergent gene expression broadly. The consequences of whole tissue sampling of complex tissues have the potential to alter not only inferred gene ontological processes, but also the structure and evolution of gene networks, the relative importance of cis- and trans-regulatory evolution, and even insights into the processes and rates underlying expression evolution.

## Acknowledgements

This work was supported by an NSF Graduate Research Fellowship to KEH (DGE-2034612), an NSF grant to ELL (DEB 1557059), and grants from the Eunice Kennedy Shriver National Institute of Child Health and Human Development of the National Institutes of Health (R01-HD073439, R01-HD094787) to JMG. We would like to thank Jonathan Velotta and members of the Larson and Tinghitella Labs for feedback on this project and Ben Fotovich, Ivan Kovanda, and the IT Department at the University of Denver for support using the High Performance Computing Cluster.

## Author contributions

KEH, ELL, and JMG conceived of the study. KEH conducted the analyses. KEH and ELL wrote the manuscript with input from JMG.

## Data accessibility

There is no data to be archived. Scripts used in the manuscript will be available upon publication at https://github.com/KelsieHunnicutt.

