## Supplemental Materials for "Unraveling patterns of disrupted gene expression across a complex tissue"

**Supplemental Figures:**

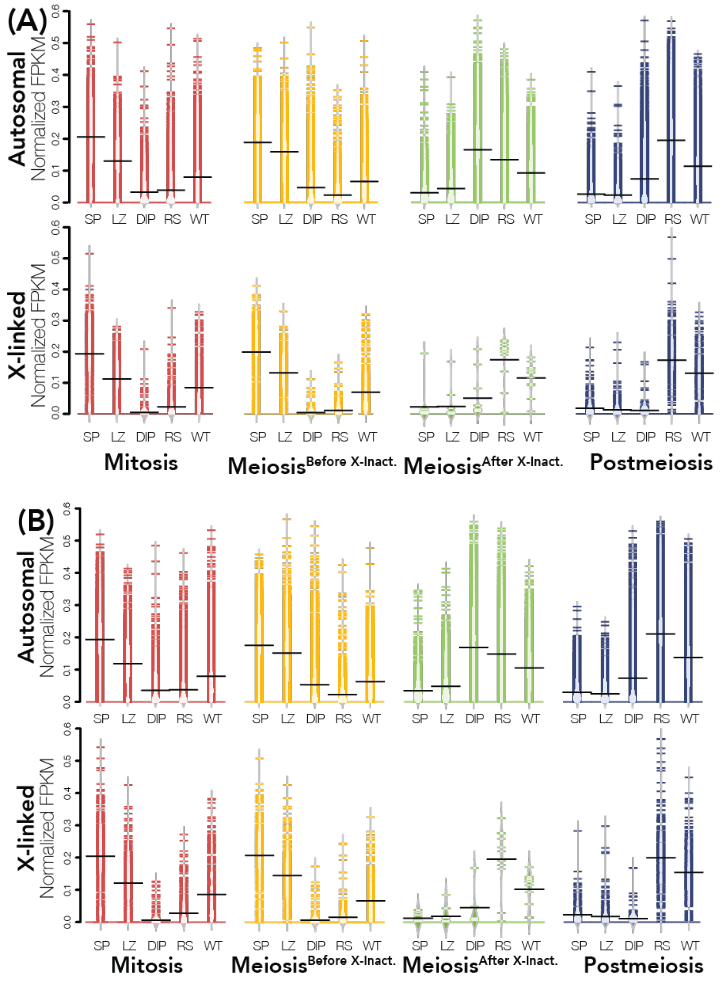


**Figure S1. Expression of induced genes in *mus* and *dom* samples.** For each sorted cell population, we defined a set of induced genes in parental samples that had a median expression two times greater than the median expression of those genes across the remaining cell types. Expression of induced genes in *mus* **(A)** and *dom* **(B)** individuals is plotted across all sample types (SP = Mitosis, LZ = Meiosis$^{\text{Before X-Inact.}}$, DIP = Meiosis$^{\text{After X-Inact.}}$, RS = Postmeiosis, and WT = Whole Testes) with cell type of induced genes indicated by color (red = Mitosis, yellow = Meiosis$^{\text{Before X-Inact.}}$, green = Meiosis$^{\text{After X-Inact.}}$, and blue = Postmeiosis). FPKM is normalized so that the sum of squares equals 1 using the R package vegan (Oksanen et al. 2007).

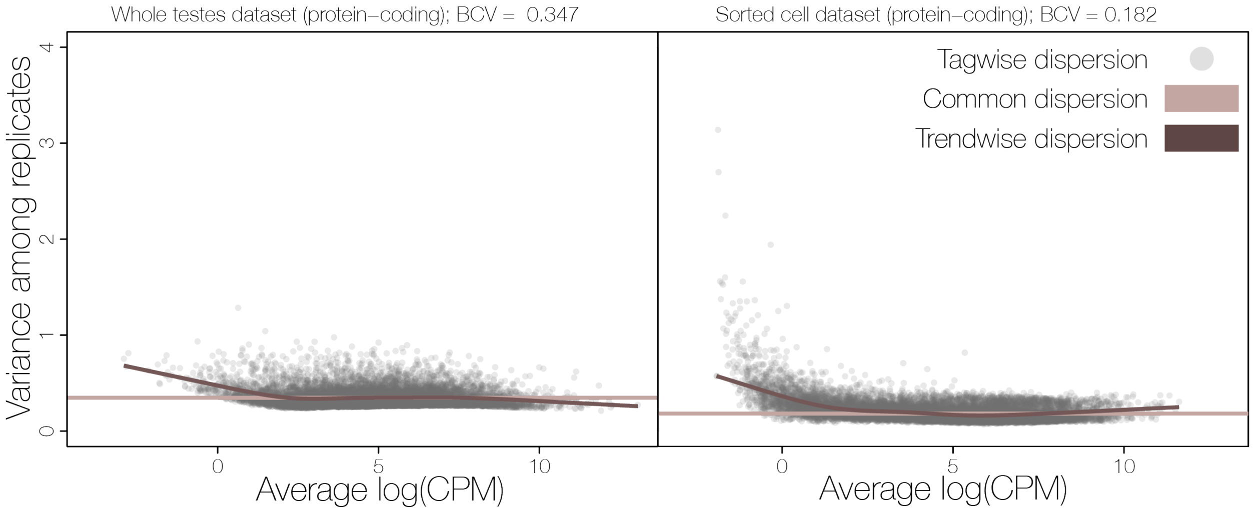


**Fig S2. Dispersion estimates and biological coefficients of variation (BCV) across protein-coding genes for the whole testes and sorted cell datasets**. All dispersion estimates were calculated in R with the edgeR package (McCarthy et al. 2012). Common dispersion for each dataset is calculated using a common estimate across all genes (taupe line). The trendwise dispersion calculation fits an estimate of dispersion based on the mean-variance trend across the entire dataset so that genes with similar abundances have similar variance estimates (brown line). Tagwise dispersion estimates dispersion on a per gene basis (gray dots). The BCV is the square root of the common dispersion.

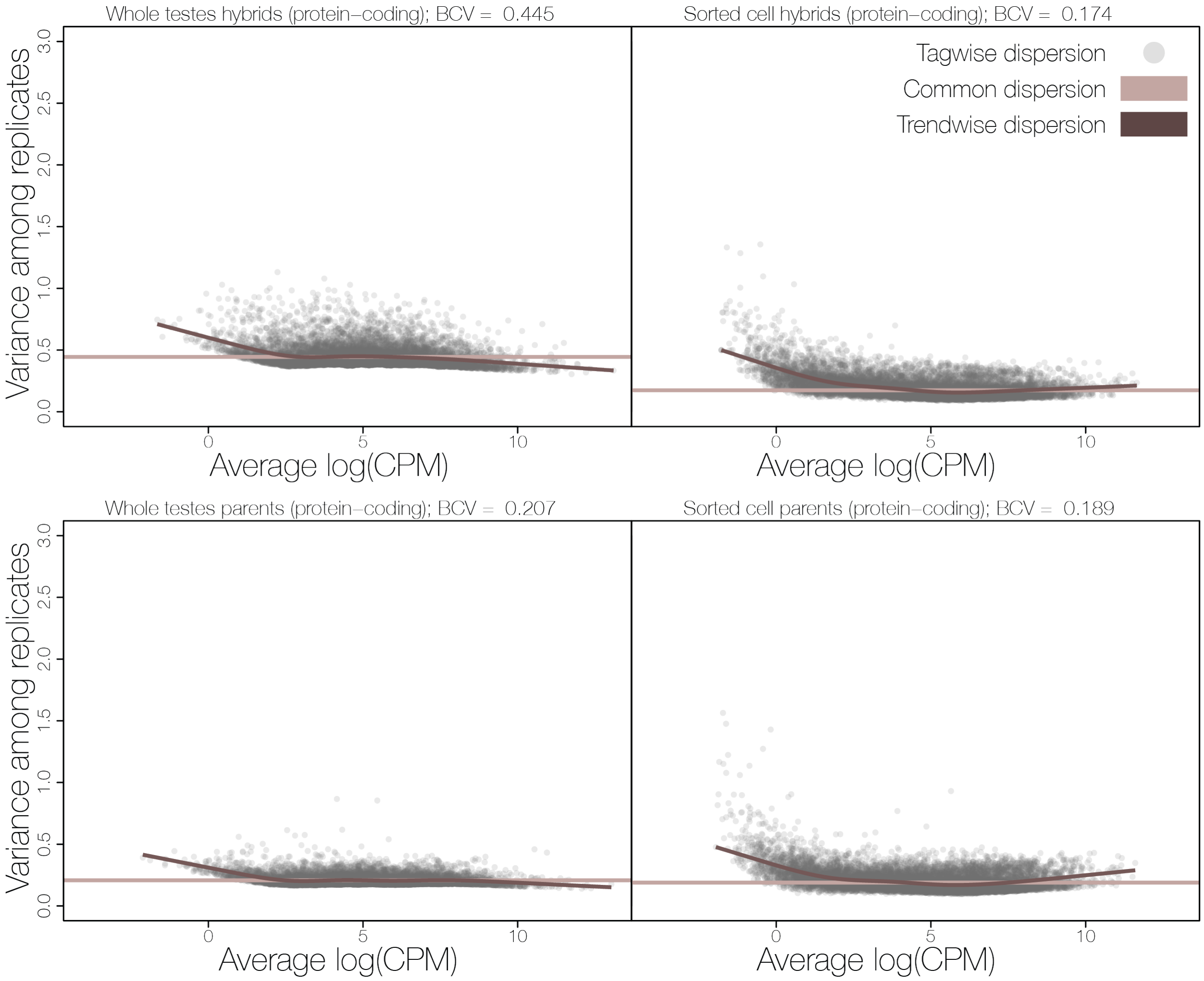


**Fig S3. Dispersion estimates and biological coefficient of variation (BCV) calculations across protein-coding genes for parental and hybrid samples separately for the whole testes and sorted cell datasets.** All dispersion estimates were calculated in R with the edgeR package (McCarthy et al. 2012). Common dispersion for each dataset is calculated using a common estimate across all genes (taupe line). The trendwise dispersion calculation fits an estimate of dispersion based on the mean-variance trend across the entire dataset so that genes with similar abundances have similar variance estimates (brown line). Tagwise dispersion estimates dispersion on a per gene basis (gray dots). The BCV is the square root of the common dispersion.

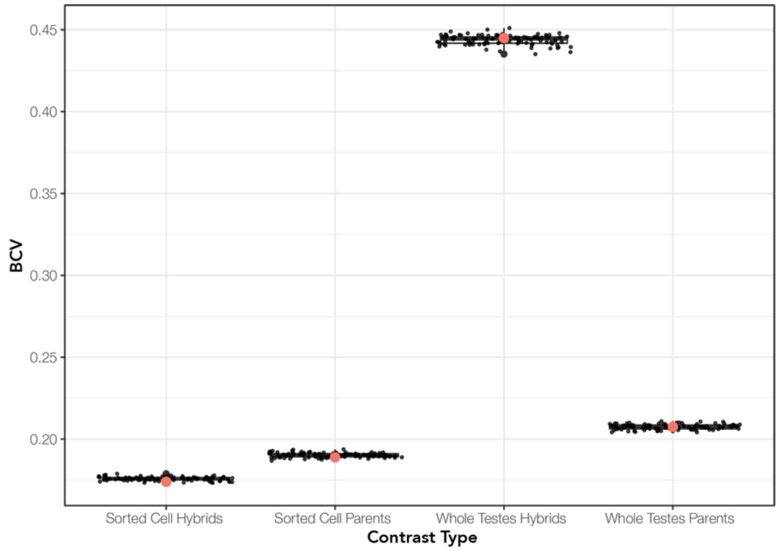


**Fig S4. BCV bootstrap estimates from the first bootstrapping approach.** We randomly sampled a set of 10000 genes for 100 replicates (bootstraps) from the raw count files generated by featureCount using only protein-coding genes then computed the BCV with the edgeR package (McCarthy et al. 2012) from these samples. Red dots indicate the BCV calculated from the full dataset.

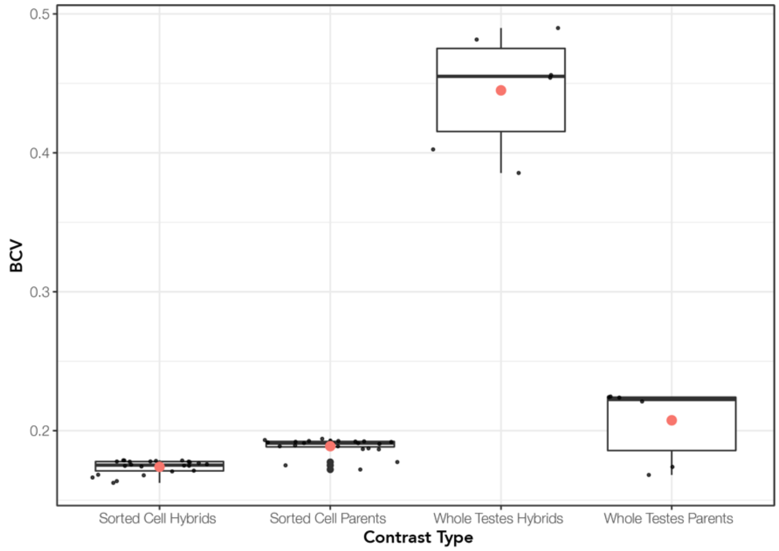


**Fig S5. BCV bootstrap estimates from the second bootstrapping approach.** We dropped one individual per contrast type then recalculated the BCV using only protein-coding genes for that contrast type across all combinations of individuals. The BCV was calculated with the edgeR package (McCarthy et al. 2012) from these samples. Red dots indicate the BCV calculated from the full dataset.

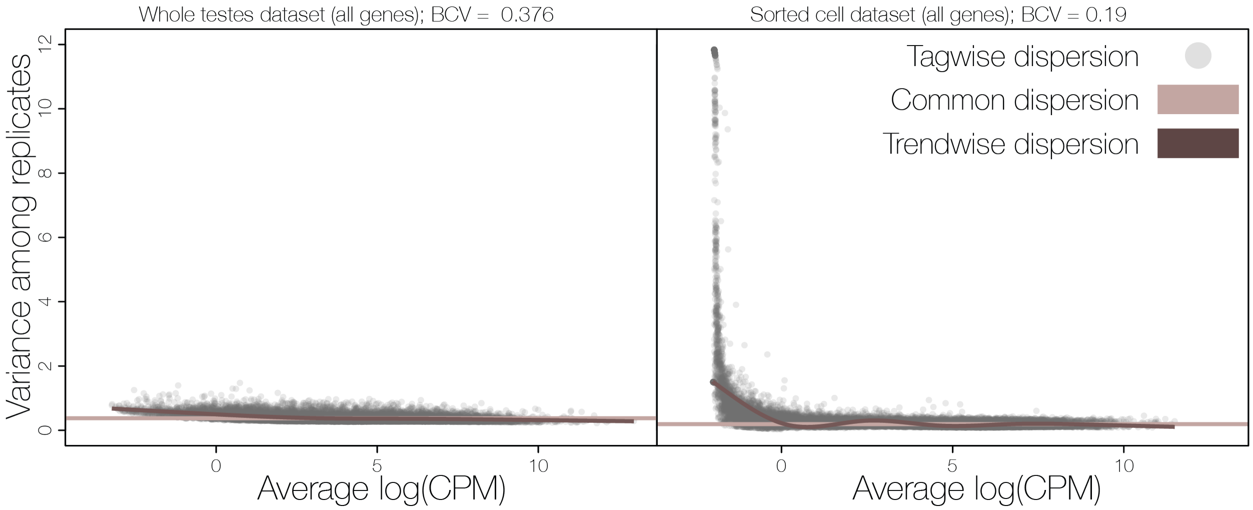


**Fig S6. Dispersion estimates and biological coefficients of variation (BCV) across all genes for the whole testes and sorted cell datasets.** All dispersion estimates were calculated in R with the edgeR package (McCarthy et al. 2012). Common dispersion for each dataset is calculated using a common estimate across all genes (taupe line). The trendwise dispersion calculation fits an estimate of dispersion based on the mean-variance trend across the entire dataset so that genes with similar abundances have similar variance estimates (brown line). Tagwise dispersion estimates dispersion on a per gene basis (gray dots). The BCV is the square root of the common dispersion.

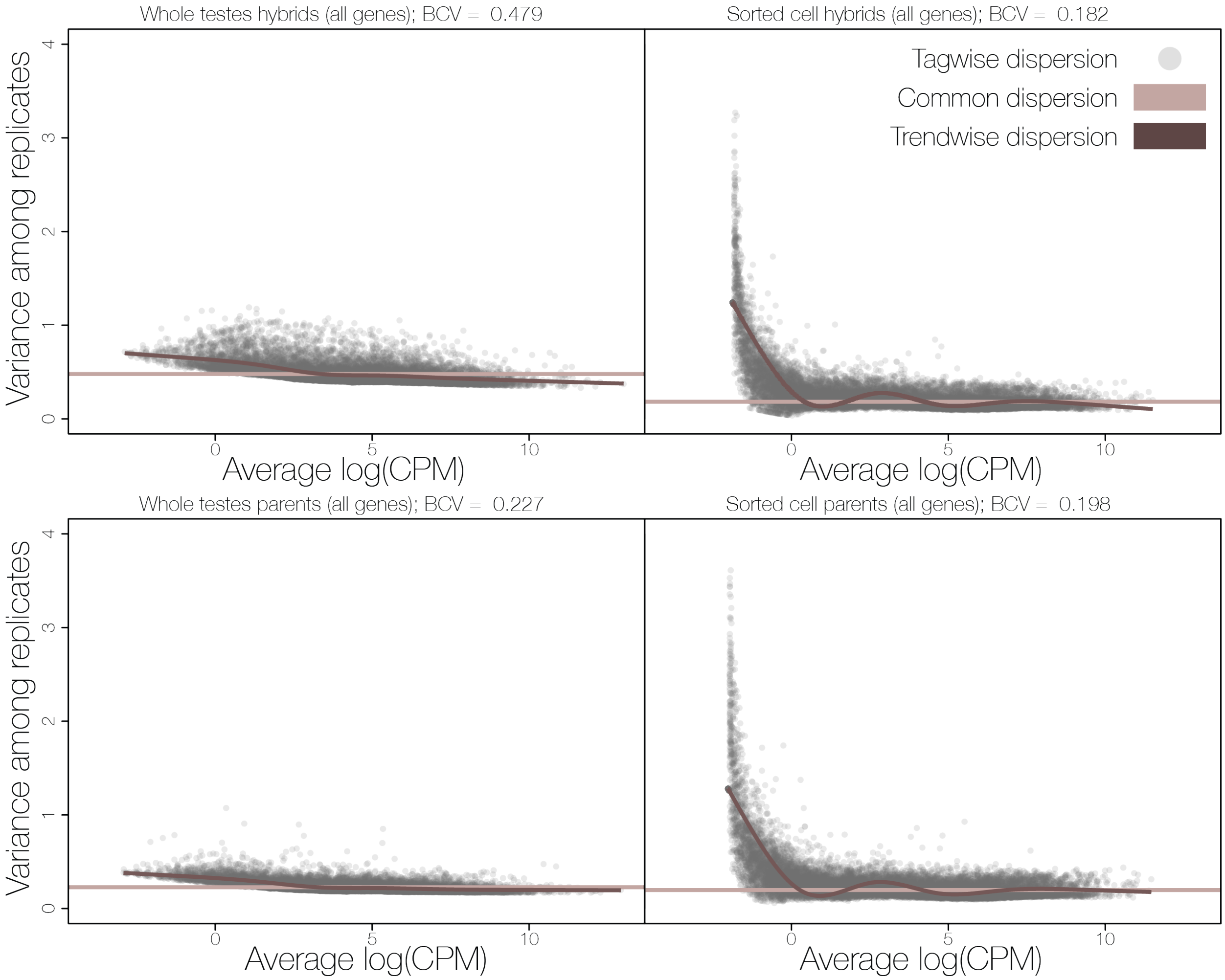


**Fig S7. Dispersion estimates and biological coefficient of variation (BCV) calculations across all genes for parental and hybrid samples separately for the whole testes and sorted cell datasets.** All dispersion estimates were calculated in R with the edgeR package (McCarthy et al. 2012). Common dispersion for each dataset is calculated using a common estimate across all genes (taupe line). The trendwise dispersion calculation fits an estimate of dispersion based on the mean-variance trend across the entire dataset so that genes with similar abundances have similar variance estimates (brown line). Tagwise dispersion estimates dispersion on a per gene basis (gray dots). The BCV is the square root of the common dispersion.

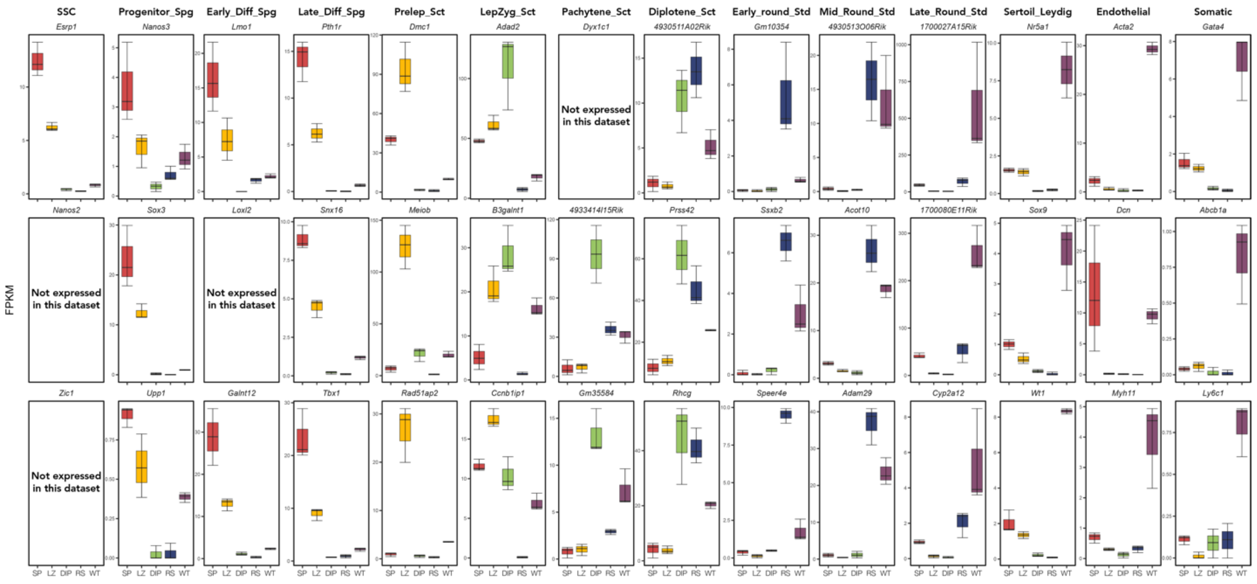


**Fig S8. *Dom* whole testes expression profiles show signatures of many diverse cell types using a second set of marker genes identified using single cell RNASeq.** Expression of cell-specific marker genes (from Hermann et al. 2018) across each sample type for *dom* reference samples. We quantified expression (FPKM) of three marker genes (rows) associated with testes-specific cell types (columns). Each panel displays marker expression in each sample type (red = Mitosis (SP), yellow = Meiosis$^{\text{Before X-Inact.}}$ (LZ), green = Meiosis$^{\text{After X-Inact.}}$ (DIP), blue = Postmeiosis (RS), and purple = Whole Testes (WT)). Marker genes correspond to the following stages: SSC - Spermatogonial stem cells, Progenitor_Spg - Progenitor spermatogonia, Early_Diff_Spg - Early differentiating spermatogonia, Late_Diff_Spg - Late differentiating spermatogonia, Prelep_Sct - Pre-leptotene spermatocytes, LepZyg_Sct - Leptotene-zygotene spermatocytes, Pachytene_Sct - Pachytene spermatocytes, Diplotene_Sct - Diplotene spermatocytes, Early_round_Std - Early round spermatids, Mid_Round_Std - Midpoint round spermatids, Late_Round_Std - Late round spermatids, Sertoil_Leydig - Sertoli and Leydig cells, Endothelial - Endothelial cells, and Somatic - Somatic cells.

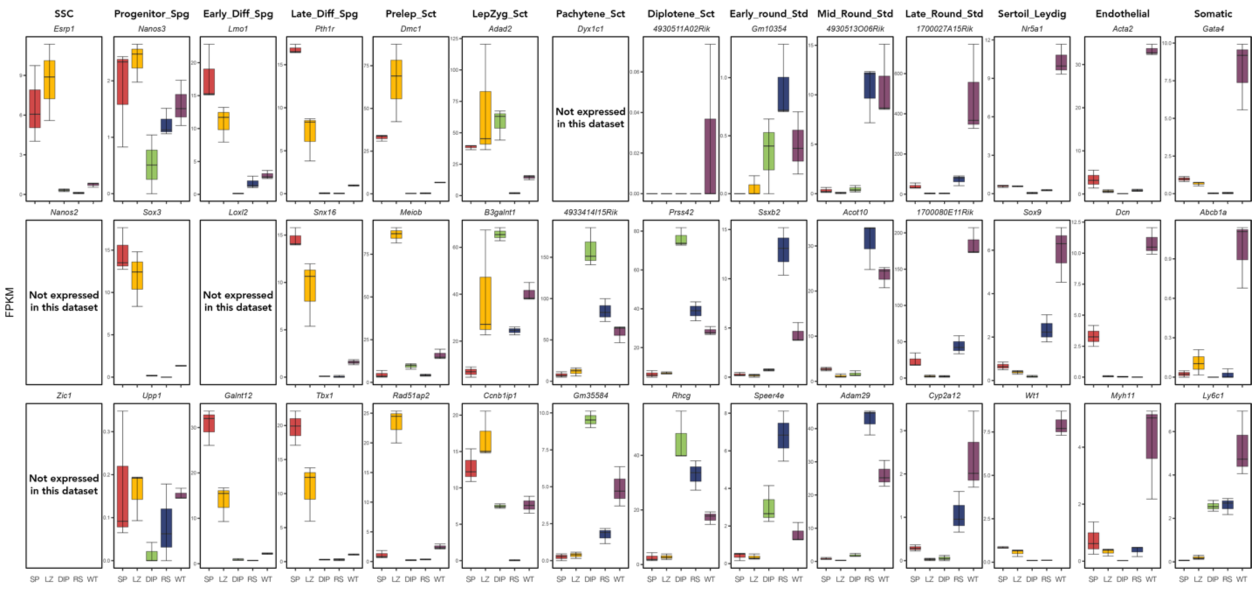


**Fig S9. *Mus* whole testes expression profiles show signatures of many diverse cell types using a second set of marker genes identified using single cell RNASeq.** Expression of cell-specific marker genes (from Hermann et al. 2018) across each sample type for *mus* reference samples. We quantified expression (FPKM) of three marker genes (rows) associated with testes-specific cell types (columns). Each panel displays marker expression in each sample type (red = Mitosis (SP), yellow = Meiosis$^{\text{Before X-Inact.}}$ (LZ), green = Meiosis$^{\text{After X-Inact.}}$ (DIP), blue = Postmeiosis (RS), and purple = Whole Testes (WT)). Marker genes correspond to the following stages: SSC - Spermatogonial stem cells, Progenitor_Spg - Progenitor spermatogonia, Early_Diff_Spg - Early differentiating spermatogonia, Late_Diff_Spg - Late differentiating spermatogonia, Prelep_Sct - Pre-leptotene spermatocytes, LepZyg_Sct - Leptotene-zygotene spermatocytes, Pachytene_Sct - Pachytene spermatocytes, Diplotene_Sct - Diplotene spermatocytes, Early_round_Std - Early round spermatids, Mid_Round_Std - Midpoint round spermatids, Late_Round_Std - Late round spermatids, Sertoil_Leydig - Sertoli and Leydig cells, Endothelial - Endothelial cells, and Somatic - Somatic cells.is specific to Meiotic$^{\text{After X-Inact.}}$ cells (Nguyen et al. 2002).

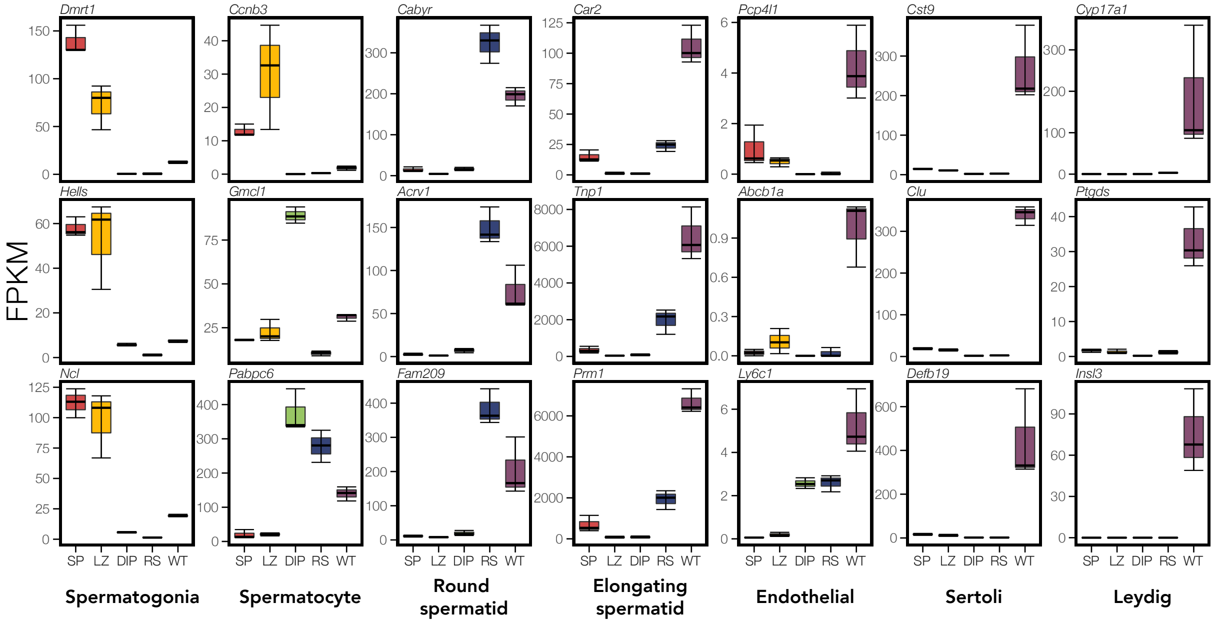


**Fig S10. *Mus* whole testes expression profiles show signatures of many diverse cell types.** Expression of cell-specific marker genes (from Green et al. 2018) across each sample type for *mus* reference samples. We quantified expression (FPKM) of three marker genes (rows) associated with testes-specific cell types (columns). Each panel displays marker expression in each sample type (red = Mitosis (SP), yellow = Meiosis$^{\text{Before X-Inact.}}$ (LZ), green = Meiosis$^{\text{After X-Inact.}}$ (DIP), blue = Postmeiosis (RS), and purple = Whole Testes (WT)). Note, *Ccnb3* expression is specific to Meiotic$^{\text{Before X-Inact.}}$ cells (Maekawa et al. 2004), and *Gmc1* is specific to Meiotic$^{\text{After X-Inact.}}$ cells (Nguyen et al. 2002).

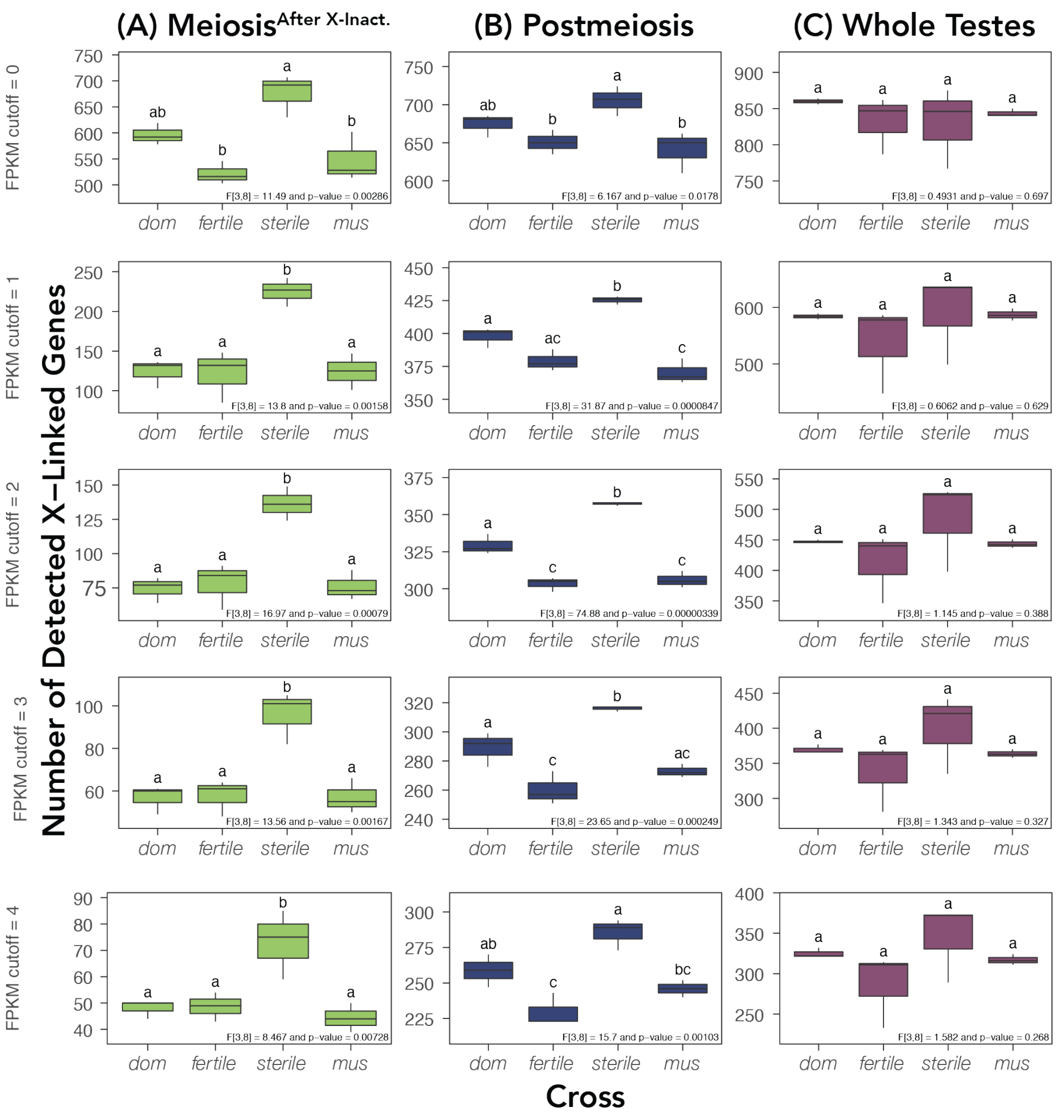


**Fig S11. Overexpression of the X chromosome in *sterile* hybrids is detectable in sorted cell populations but not whole testes.** Mean number of X-linked genes for each cross within a sample type with a minimum expression of the indicated FPKM along the left side of the panel across all samples within a dataset (sorted cells or whole testes). Whiskers show 95% confidence intervals. F-statistics and p-values from each one-way ANOVA are presented in the bottom right of each panel. Different letters above error bars indicate a significant difference between means at $p<0.05$ using a post-hoc Tukey HSD test. Each column shows results from each sample type where X overexpression is expected in *sterile* hybrids compared to parental mice with the same X chromosome (*i.e., mus*), Meiosis$^{\text{After X-Inact.}}$ (**A**; green), Postmeiosis (**B**; blue), and Whole Testes (**C**; purple).

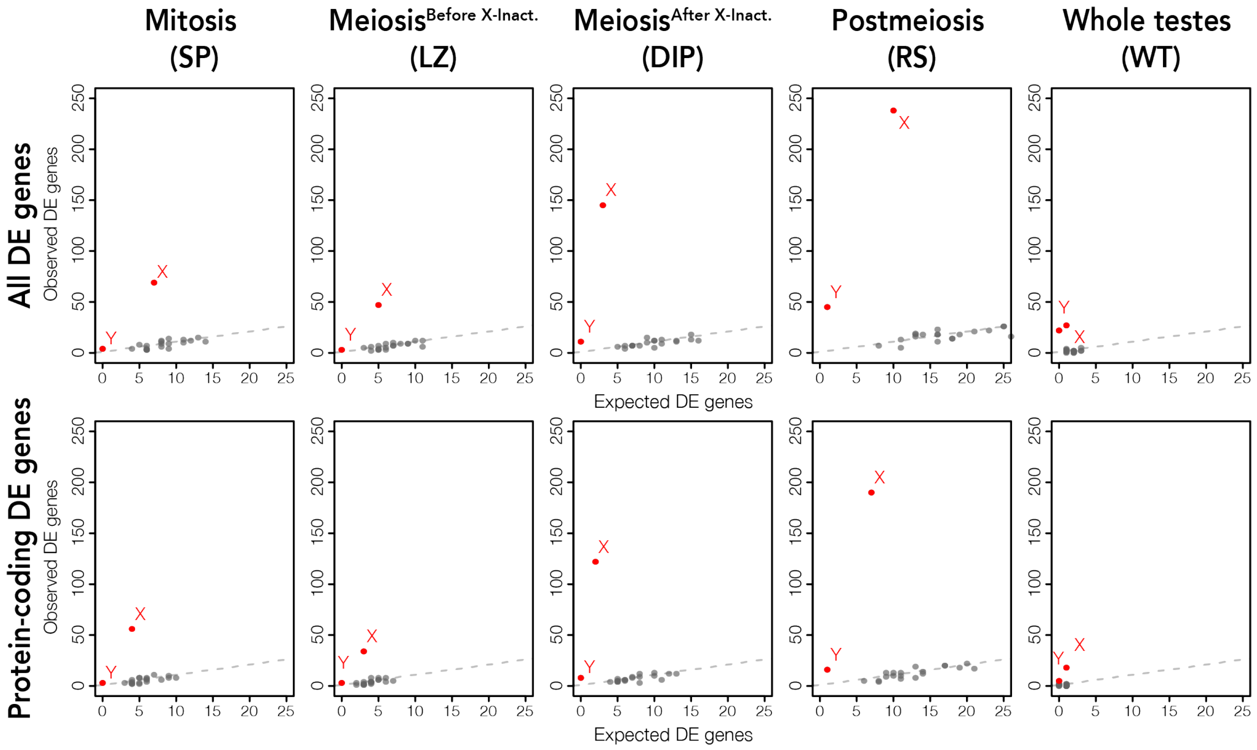


**Fig S12. Sex chromosomes are enriched for DE genes across all stages.** For each sample type, a scatter plot displays expected versus observed counts of DE genes for each chromosome. Chromosomes above the dashed line are over-enriched for DE genes, and chromosomes below the dashed line are under-enriched for DE genes, with chromosomes were p-values were less than 0.001 after FDR correction are highlighted in red and labelled. Upper panels are DE genes from all annotated genes and lower panels are DE genes from only protein-coding genes.

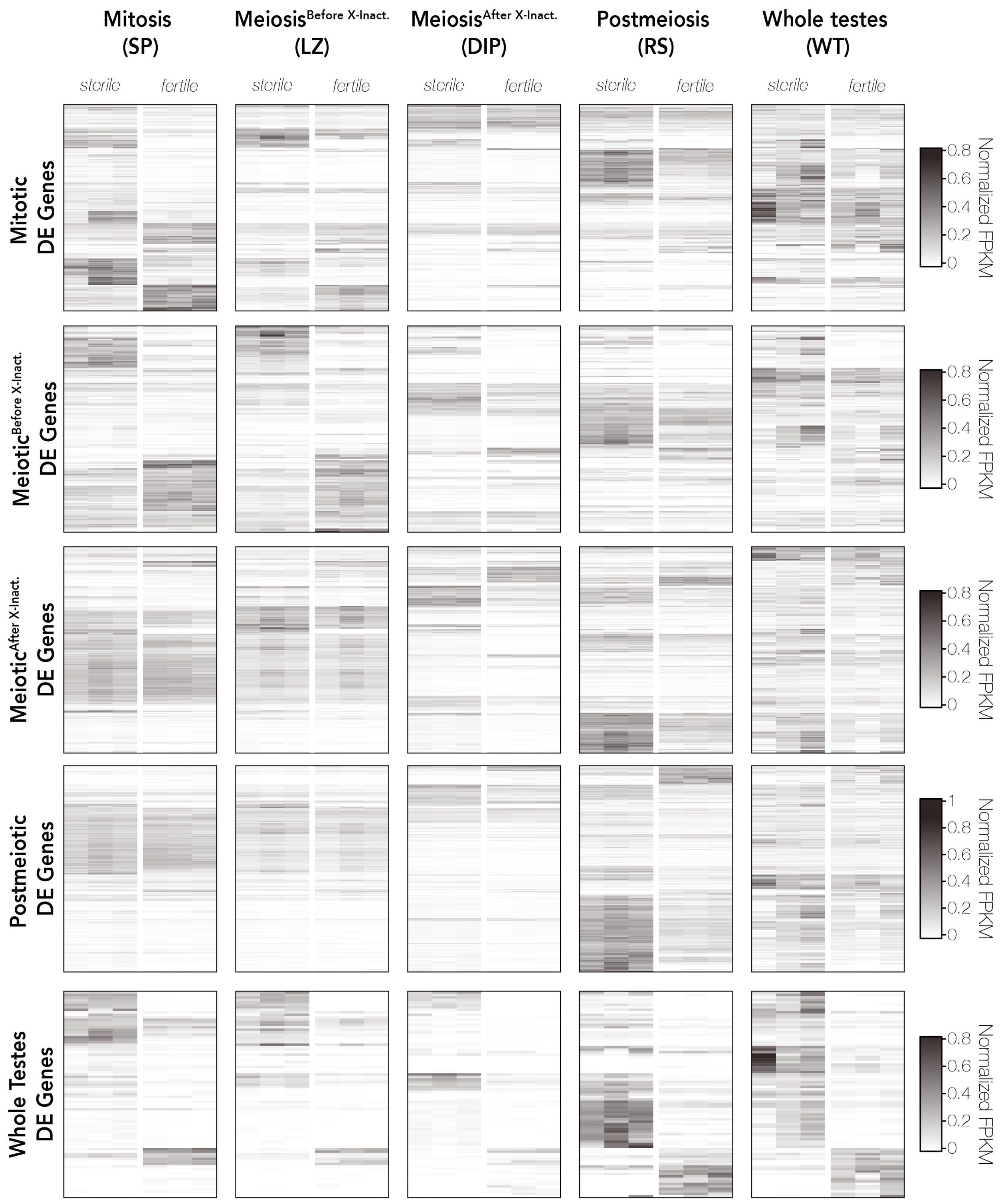


**Fig S13. Expression of sample-type specific DE genes in *sterile* and *fertile* hybrids across all sample types.** FPKM values were normalized so that the sum of squares equals one using the R package vegan (Oksanen et al. 2007). Beanplots were generated with the R package beanplot (Kampstra 2008). Each heatmap has gene expression plotted as normalized FPKM values that are hierarchically clustered using Euclidean distance. Each row plots expression across one gene and darker colors indicate higher expression. Heatmaps were generated with the R package ComplexHeatmap v.2.3.2 (Gu et al. 2016).

**Supplemental Tables:**

Table S1: Sample information and read counts. Sample IDs correspond to the cross of the individual (CC = CZECHII/EiJ, LL = LEWES/EiJ, PP = PWK/PhJ, and WW = WSB/EiJ), the individuals ID number, and the sample type (SP = Mitosis, LZ = Meiosis$^{\text{Before X-Inact.}}$, DIP = Meiosis$^{\text{After X-Inact.}}$, RS = Postmeiosis, and WT = Whole Testes).

| Sample ID | SRA Accession | Raw Reads (F only) | Post-Tophat PWK-Alignment Reads (F+R) | Post-Tophat WSB-Alignment Reads (F+R) | Post-Suspenders Reads (F+R) | Assigned FeatureCount Read Pairs (no multi-mapped reads) |
|---|---|---|---|---|---|---|
| CCPP-21.1-DIP | SRR2761570, SRR2761592 | 21477056 | 26340755 | 25674973 | 19232543 | 7381118 |
| CCPP-21.1-LZ | SRR2761571, SRR2761593 | 26998246 | 33134843 | 32416769 | 23610512 | 8867722 |
| CCPP-21.1-RS | SRR2761572, SRR2761594 | 22195796 | 35312787 | 34717874 | 19809616 | 7721308 |
| CCPP-21.1-SP | SRR2761573, SRR2761595 | 31751428 | 38868844 | 37967473 | 27275613 | 9931488 |
| CCPP-21.2-DIP | SRR2761596 | 36852964 | 46965915 | 45728821 | 33544699 | 12803354 |
| CCPP-21.2-LZ | SRR2761574, SRR2761597 | 26549896 | 32060131 | 31287872 | 23526641 | 8786313 |
| CCPP-21.2-RS | SRR2761598 | 22235268 | 36029725 | 35270117 | 20281803 | 7942450 |
| CCPP-21.2-SP | SRR2761575, SRR2761599 | 28802542 | 34567190 | 33765425 | 24615826 | 8998231 |
| CCPP-21.3-DIP | SRR2761549, SRR2761600 | 22468784 | 27090066 | 26309504 | 20152293 | 7764490 |
| CCPP-21.3-LZ | SRR2761550, SRR2761601 | 20428992 | 24566552 | 23933367 | 17891667 | 6709337 |
| CCPP-21.3-RS | SRR2761602 | 32112934 | 45597240 | 44284041 | 28554464 | 11260227 |
| CCPP-21.3-SP | SRR2761551, SRR2761603 | 21751826 | 26924883 | 26217413 | 19057748 | 7075581 |
| LLPP-17.2-DIP | SRR2761576, SRR2761604 | 19989428 | 24880530 | 24872825 | 18021849 | 6822392 |
| LLPP-17.2-RS | SRR2761577, SRR2761605 | 22525160 | 33916646 | 33841987 | 20044435 | 7762604 |
| LLPP-18.1-LZ | SRR2761578, SRR2761606 | 27817084 | 35876214 | 35985195 | 24854243 | 8985489 |
| LLPP-19.1-DIP | SRR2761552, SRR2761607 | 18058676 | 22479413 | 22438712 | 16204722 | 6096994 |
| LLPP-19.1-SP | SRR2761579, SRR2761608 | 33430222 | 39885520 | 39708715 | 28440056 | 10028665 |
| LLPP-19.2-RS | SRR2761553, SRR2761609 | 18407584 | 28601605 | 28594856 | 16484648 | 6323895 |
| LLPP-19.3-DIP | SRR2761554, SRR2761610 | 19415690 | 23093669 | 23054030 | 17375360 | 6661074 |
| LLPP-19.3-RS | SRR2761555, SRR2761611 | 19781658 | 27402180 | 27494017 | 17681421 | 6959808 |
| LLPP-22.7-LZ | SRR2761556, SRR2761612 | 20036778 | 25245447 | 25262290 | 17820330 | 6477961 |
| LLPP-22.7-SP | SRR2761557, SRR2761613 | 18338020 | 22593735 | 22567117 | 16094821 | 5971162 |
| LLPP-22.8-LZ | SRR2761558, SRR2761614 | 23411888 | 29264006 | 29336299 | 21033663 | 7685705 |
| LLPP-22.8-SP | SRR2761559, SRR2761615 | 19297400 | 22586043 | 22519133 | 16561826 | 6017707 |
| LLPP-272-WT | SRR2060953 | 124219124 | 142525063 | 142380020 | 108427854 | 42898717 |
| LLPP-290-WT | SRR2060952 | 34464432 | 38526459 | 38543523 | 32163922 | 13313386 |
| LLPP-93-WT | SRR2060950 | 63070002 | 47005603 | 47097532 | 34088411 | 12586942 |
| LLWW-148-WT | SRR2060837 | 101248844 | 74722027 | 77167079 | 54406591 | 19891163 |
| LLWW-149-WT | SRR2060842 | 107432468 | 128083415 | 130793609 | 93850300 | 35897894 |
| LLWW-150-WT | SRR2060843 | 100497198 | 108902509 | 111315069 | 81179922 | 31309028 |

| | | | | | | |
|---|---|---|---|---|---|---|
| PPCC-151-WT | SRR2060844 | 76325172 | 58327482 | 56337894 | 39979615 | 14783396 |
| PPCC-152-WT | SRR2060846 | 98804204 | 118179509 | 115163595 | 85840471 | 33494033 |
| PPCC-170-WT | SRR2060939 | 76594232 | 90758054 | 88360116 | 66410204 | 25980058 |
| PPLL-131-WT | SRR2060954 | 24882156 | 28222294 | 28244608 | 23110713 | 9474627 |
| PPLL-15.2-DIP | SRR2761580, SRR2761616 | 23647380 | 30142023 | 29984831 | 21268666 | 8069338 |
| PPLL-15.2-RS | SRR2761581, SRR2761617 | 23128976 | 40531135 | 40317066 | 20658116 | 7842258 |
| PPLL-16.1-DIP | SRR2761560, SRR2761618 | 20048656 | 24991779 | 24867469 | 17763716 | 6646884 |
| PPLL-16.1-LZ | SRR2761561, SRR2761619 | 21281980 | 27273010 | 27117068 | 18786051 | 6912012 |
| PPLL-16.1-RS | SRR2761562, SRR2761620 | 19168394 | 29754797 | 29751712 | 16517064 | 6361843 |
| PPLL-16.1-SP | SRR2761563, SRR2761621 | 22451632 | 27796017 | 27662178 | 19388612 | 7051214 |
| PPLL-17.1-DIP | SRR2761622 | 37268646 | 45864903 | 45637800 | 33049299 | 12492601 |
| PPLL-17.1-LZ | SRR2761582, SRR2761623 | 30492632 | 41043284 | 40790863 | 27195859 | 9923511 |
| PPLL-17.1-RS | SRR2761624 | 25277730 | 40651852 | 40533661 | 22361132 | 8609292 |
| PPLL-17.1-SP | SRR2761583, SRR2761625 | 33687062 | 40791246 | 40560457 | 29439375 | 11005974 |
| PPLL-17.3-LZ | SRR2761564, SRR2761626 | 19742116 | 25086723 | 24944022 | 17398240 | 6412712 |
| PPLL-17.3-SP | SRR2761565, SRR2761627 | 23598202 | 28711605 | 28465007 | 20271530 | 7299012 |
| PPLL-278-WT | SRR2060955 | 102188034 | 133331212 | 133813859 | 89775343 | 33934908 |
| PPLL-52-WT | SRR2060951 | 44950474 | 53997983 | 53670612 | 35822322 | 12838352 |
| WWLL-3.1-DIP | SRR2761584, SRR2761628 | 21970318 | 26565246 | 27212060 | 19982585 | 7588358 |
| WWLL-3.1-RS | SRR2761585, SRR2761629 | 22523002 | 33735184 | 34378293 | 20360830 | 7972083 |
| WWLL-3.1-SP | SRR2761566,SRR2761630 | 21104146 | 25684138 | 26185875 | 18283388 | 6458669 |
| WWLL-4.1-LZ | SRR2761567, SRR2761631 | 20075246 | 25176105 | 25778595 | 17983406 | 6642792 |
| WWLL-6.1-RS | SRR2761568, SRR2761632 | 20388762 | 29125360 | 29720516 | 18060410 | 7053241 |
| WWLL-7.1-DIP | SRR2761569, SRR2761633 | 19974584 | 23300740 | 24024169 | 17794108 | 6798887 |
| WWLL-7.2-DIP | SRR2761586, SRR2761634 | 20182808 | 24558522 | 25232730 | 18511737 | 6969049 |
| WWLL-7.2-LZ | SRR2761587, SRR2761635 | 20087232 | 25629600 | 26319298 | 18253476 | 6675109 |
| WWLL-7.2-RS | SRR2761588, SRR2761636 | 20822386 | 28662881 | 29206953 | 18976974 | 7607417 |
| WWLL-7.2-SP | SRR2761589, SRR2761637 | 25852196 | 32343831 | 33080949 | 22904315 | 8266300 |
| WWLL-7.3-LZ | SRR2761590, SRR2761638 | 41266036 | 53274705 | 54449278 | 37203850 | 13432758 |
| WWLL-7.3-SP | SRR2761591, SRR2761639 | 35512824 | 43398931 | 44340114 | 31279482 | 11230973 |

Table S2: Counts of different categories of hybrid DE genes for each sample type.

| Stage | X-linked DE genes | Y-linked DE genes | Autosomal DE genes | Up-regulated in *sterile* | Down-regulated in *sterile* | Total DE genes |
|---|---|---|---|---|---|---|
| Mitosis | 69 | 4 | 158 | 152 | 79 | 231 |
| Meiosis$^{\text{Before X-Inact.}}$ | 47 | 3 | 128 | 88 | 90 | 178 |
| Meiosis$^{\text{After X-Inact.}}$ | 145 | 11 | 187 | 284 | 59 | 343 |
| Postmeiosis | 238 | 45 | 323 | 497 | 109 | 606 |
| Whole Testes | 27 | 22 | 34 | 63 | 20 | 83 |

Table S3: The number of observed and expected number of X-linked DE genes and the significance of the hypergeometric test for enrichment of the X chromosome for DE genes for each sample type.

| Sample Type | X-linked Expected DE Genes | X-linked Observed DE Genes | P-Value |
|---|---|---|---|
| Mitosis | 7 | 69 | 0 |
| Meiosis$^{\text{Before X-Inact.}}$ | 5 | 47 | 0 |
| Meiosis$^{\text{Before X-Inact.}}$ | 3 | 145 | 0 |
| Postmeiosis | 10 | 238 | 0 |
| Whole Testes | 1 | 27 | 0 |

Table S4: The number of observed and expected number of Y-linked DE genes and the significance of the hypergeometric test for enrichment of the Y chromosome for DE genes for each sample type.

| Sample Type | Y-linked Expected DE Genes | Y-linked Observed DE Genes | P-Value |
|---|---|---|---|
| Mitosis | 0 | 4 | 0 |
| Meiosis$^{\text{Before X-Inact.}}$ | 0 | 3 | 0 |
| Meiosis$^{\text{After X-Inact.}}$ | 0 | 11 | 0 |
| Postmeiosis | 1 | 45 | 0 |
| Whole Testes | 0 | 22 | 0 |

Table S5: Direction of regulation (relative to *sterile*) for hybrid DE genes in whole testes and each sorted cell populations for each pairwise comparison.

| Comparison | Up in WT & down in sorted cell population | Down in WT & up in sorted cell population | Up-regulated in both | Down-regulated in both | Misregulated between sorted cell population and whole testes (%) |
|---|---|---|---|---|---|
| Mitosis and WT | 0 | 0 | 25 | 6 | 0 |
| Meiosis$^{\text{Before X-Inact.}}$ and WT | 0 | 0 | 22 | 5 | 0 |
| Meiosis$^{\text{After X-Inact.}}$ and WT | 0 | 0 | 28 | 5 | 0 |
| Postmeiosis and WT | 0 | 0 | 47 | 17 | 0 |

Table S6: Direction of regulation (relative to *mus*) for DE genes between *dom* and *mus* in whole testes and each sorted cell populations for each pairwise comparison.

| Comparison | Up in WT & down in sorted cell population | Down in WT & up in sorted cell population | Up-regulated in both | Down-regulated in both | Misregulated between sorted cell population and whole testes (%) |
|---|---|---|---|---|---|
| Mitosis and WT | 22 | 14 | 392 | 712 | 3.26 |
| Meiosis$^{\text{Before X-Inact.}}$ and WT | 10 | 15 | 357 | 553 | 2.75 |
| Meiosis$^{\text{After X-Inact.}}$ and WT | 11 | 14 | 557 | 757 | 1.9 |
| Postmeiosis and WT | 5 | 3 | 721 | 1130 | 0.43 |

Table S7: The proportion of DE genes shared between each sample type comparison (*i.e.,* # DE genes in common between sample types/# of unique DE genes in both sample types) across different log(x) Fold Change cutoffs.

| Comparison | log(0) | log(1) | log(2) | log(3) |
|---|---|---|---|---|
| Whole Testes vs. Mitosis | 0.11 | 0.182 | 0.183 | 0.236 |
| Whole Testes vs. Meiosis$^{\text{Before X-Inact.}}$ | 0.115 | 0.17 | 0.183 | 0.218 |
| Whole Testes vs. Meiosis$^{\text{After X-Inact.}}$ | 0.084 | 0.178 | 0.192 | 0.213 |
| Whole Testes vs. Postmeiosis | 0.102 | 0.217 | 0.27 | 0.261 |
| Mitosis vs. Meiosis$^{\text{Before X-Inact.}}$ | 0.236 | 0.271 | 0.328 | 0.527 |
| Mitosis vs. Meiosis$^{\text{After X-Inact.}}$ | 0.117 | 0.129 | 0.195 | 0.261 |
| Mitosis vs. Postmeiosis | 0.096 | 0.113 | 0.152 | 0.179 |
| Meiosis$^{\text{Before X-Inact.}}$ vs. Meiosis$^{\text{After X-Inact.}}$ | 0.118 | 0.143 | 0.221 | 0.279 |
| Meiosis$^{\text{Before X-Inact.}}$ vs. Postmeiosis | 0.087 | 0.105 | 0.12 | 0.167 |
| Meiosis$^{\text{After X-Inact.}}$ vs. Postmeiosis | 0.242 | 0.244 | 0.184 | 0.183 |
